# Internalizing-peptide engineering converts blocking anti-EGFR antibodies into EGFR lysosomal targeting degraders

**DOI:** 10.64898/2026.11.20.768013

**Authors:** Petar D. Petrov, Rodrigo Vinueza-Gavilanes, Antía Pazó Vicente, Manpreet Kaur, Pilar Simón-Extremera, Julieta Carabelli, Laura Padilla, Luis M. Montuenga, Alfonso Calvo, Juan Sebastián Ruiz

## Abstract

Conventional antibodies targeting the epidermal growth factor receptor (EGFR) primarily inhibit receptor activation rather than efficiently eliminating the receptor from tumor cells. Here, we investigated whether engineering full-length anti-EGFR antibodies with internalizing peptides could promote receptor degradation, enhance their biological activity and enable the development of recombinant receptor-depleting antibodies.

Chemical conjugation of cetuximab (CTX) to the polyarginine peptide R12 enhanced intracellular uptake, reduced surface-accessible EGFR and decreased total receptor abundance *in vitro*. CTX-R12 conjugate also enhanced inhibition of wound repopulation in non-small cell lung cancer (NSCLC) cell lines, an effect reproduced using the R12 variant LINC18, and transferred to the independent blocking anti-EGFR antibody LEI. Incorporation of LINC19, a variant of LINC18, into recombinant full-length antibody fusion proteins preserved their activity. The LEI-LINC19 fusion (LEIP), reduced surface-accessible and total EGFR, with bafilomycin A1 reversing receptor loss and supporting an acidification-dependent endolysosomal component of degradation.

Functional studies indicated that the antibody-peptide reduced DNA synthesis rate. Furthermore, siRNA-mediated EGFR depletion increased TKI-associated reductions in cellular biomass in PC9 cells and increased gefitinib responsiveness in H1975 cells. Consistently, combining LEIP with either osimertinib or gefitinib produced greater inhibition of H1975 wound repopulation than the corresponding individual treatments. LDH analysis revealed increased membrane-damage-associated LDH release following LEIP-gefitinib treatment but no additional increase with LEIP-osimertinib relative to osimertinib alone.

These effects extended to 3D NSCLC models. CTX-R12 inhibited PC9 3D colony development, whereas LEIP restricted expansion of established PC9 colonies and reduced the projected area of H1975 spheroids. In H1975 spheroids initiated with 1,000 or 5,000 cells per well, LEIP-containing combinations with osimertinib or gefitinib produced greater inhibition of the spheroid size than the corresponding TKI-only treatments. Nuclear Green DCS1 staining revealed no detectable baseline core necrosis in untreated H1975 spheroids, whereas LEIP treatment was associated with DNA-positive material within and surrounding the spheroids. Moreover, LEIP-treated H1975 spheroids exhibited markedly reduced EGFR immunofluorescence compared with control and parental LEI-treated spheroids.

Together, these findings demonstrate that internalizing-peptide engineering of blocking anti-EGFR antibodies can generate full-length EGFR-depleting fusion proteins with enhanced biological activity. The recombinant LEIP format and its activity in 3D *in vitro* NSCLC tumor models support further investigation of fusion antibody-mediated EGFR degradation as a complementary strategy to EGFR kinase inhibition.

## Introduction

The epidermal growth factor receptor (EGFR) is a receptor tyrosine kinase involved in the regulation of cell proliferation, survival, differentiation and tissue organization. Aberrant EGFR signaling contributes to the development and progression of several epithelial malignancies, including non-small-cell lung cancer (NSCLC). Activating EGFR mutations define subsets of NSCLC that are particularly sensitive to small-molecule EGFR tyrosine kinase inhibitors (TKIs). Nevertheless, sustained tumor dependence on EGFR and the emergence of resistance to pharmacological kinase inhibition remain important therapeutic challenges.

Current EGFR-targeted treatments predominantly inhibit receptor function rather than eliminate the receptor protein. Cetuximab (CTX), a clinically established monoclonal antibody, binds extracellular domain III of EGFR and prevents productive ligand engagement and receptor activation [1]. However, inhibition of receptor activity and reduction of receptor abundance represent distinct biological interventions. Jaramillo et al. demonstrated that the cetuximab precursor mAb 225 undergoes EGFR-mediated internalization but that internalized antibody-receptor complexes can recycle efficiently to the plasma membrane, limiting receptor downregulation [2]. Other approaches, including combinations of antibodies recognizing noncompetitive EGFR epitopes, have been shown to interfere with receptor recycling and promote EGFR downregulation [3].

The ability to redirect cell-surface proteins toward intracellular degradation has attracted substantial interest as an alternative to conventional target inhibition. Lysosome-targeting chimeras (LYTACs) recruit endogenous lysosomal trafficking receptors to promote degradation of extracellular or membrane-associated targets [6]. Subsequently, genetically encoded cytokine receptor-targeting chimeras (KineTACs) demonstrated that recruitment of internalizing cell-surface receptors can be exploited to eliminate EGFR and other extracellular proteins [7]. These approaches establish the feasibility of receptor-directed degradation but generally require an additional trafficking-receptor recognition moiety in the degrader.

An alternative strategy is to incorporate an internalization-driving element directly into the target-binding antibody. Van Lith et al. demonstrated that conjugation of a cell-penetrating peptide to the anti-EGFR single-domain antibody 7D12 promoted receptor internalization and enhanced inhibition of EGFR activation [4]. Furthermore, Gaston et al. showed that cell-penetrating peptides can be genetically incorporated into full-length antibodies while retaining relevant antibody properties [5]. These observations raise the possibility that direct internalizing-peptide conjugation to anti-EGFR antibodies could alter their intracellular fate without deliberately recruiting a second trafficking receptor.

Chemical conjugation provides a flexible approach for evaluating internalizing peptides; however, conventional lysine-based conjugation can generate heterogeneous antibody populations with variable attachment sites and peptide-to-antibody ratios [19]. In contrast, genetic fusion enables the incorporation of internalizing peptides at predefined positions within the antibody sequence, providing controlled peptide stoichiometry at the sequence level and allowing production through established recombinant antibody-expression workflows [5].

Here, we investigated whether internalizing-peptide engineering could convert blocking anti-EGFR antibodies into receptor-depleting molecules and enhance their biological activity. We initially used chemical conjugation to evaluate R12 and its variant LINC18 in the CTX scaffold, followed by experiments using the proprietary EGFR blocking antibody LEI. We then generated recombinant CTX-LINC19 and LEI-LINC19 fusion proteins to determine whether the activity of chemically engineered antibodies could be preserved in a defined recombinant format. Surface EGFR immunofluorescence, total EGFR immunoblotting, bafilomycin A1 rescue, DNA-synthesis and wound-repopulation assays, siRNA-mediated EGFR depletion, TKI combinations and three-dimensional *in vitro* tumor models were used to investigate the mechanistic and functional consequences of this approach.

## Results

### 1. Chemical conjugation of internalizing peptides promotes cetuximab uptake and EGFR depletion

We first investigated whether chemical attachment of an internalizing peptide could alter the intracellular trafficking and receptor-depleting activity of CTX. The arginine-rich internalizing peptide R12, or its HiBiT-tagged derivative R12-HiBiT, were chemically conjugated to CTX (schematically represented in Fig. 1a, left), and intracellular antibody accumulation was evaluated in EGFR-expressing cells. Unless otherwise indicated, all antibodies were used at a concentration of 100 nM. The internalizing capacity of each peptide was previously confirmed following conjugation to an unrelated human IgG antibody (bevacizumab; see Materials and Methods).

**Figure 1.**
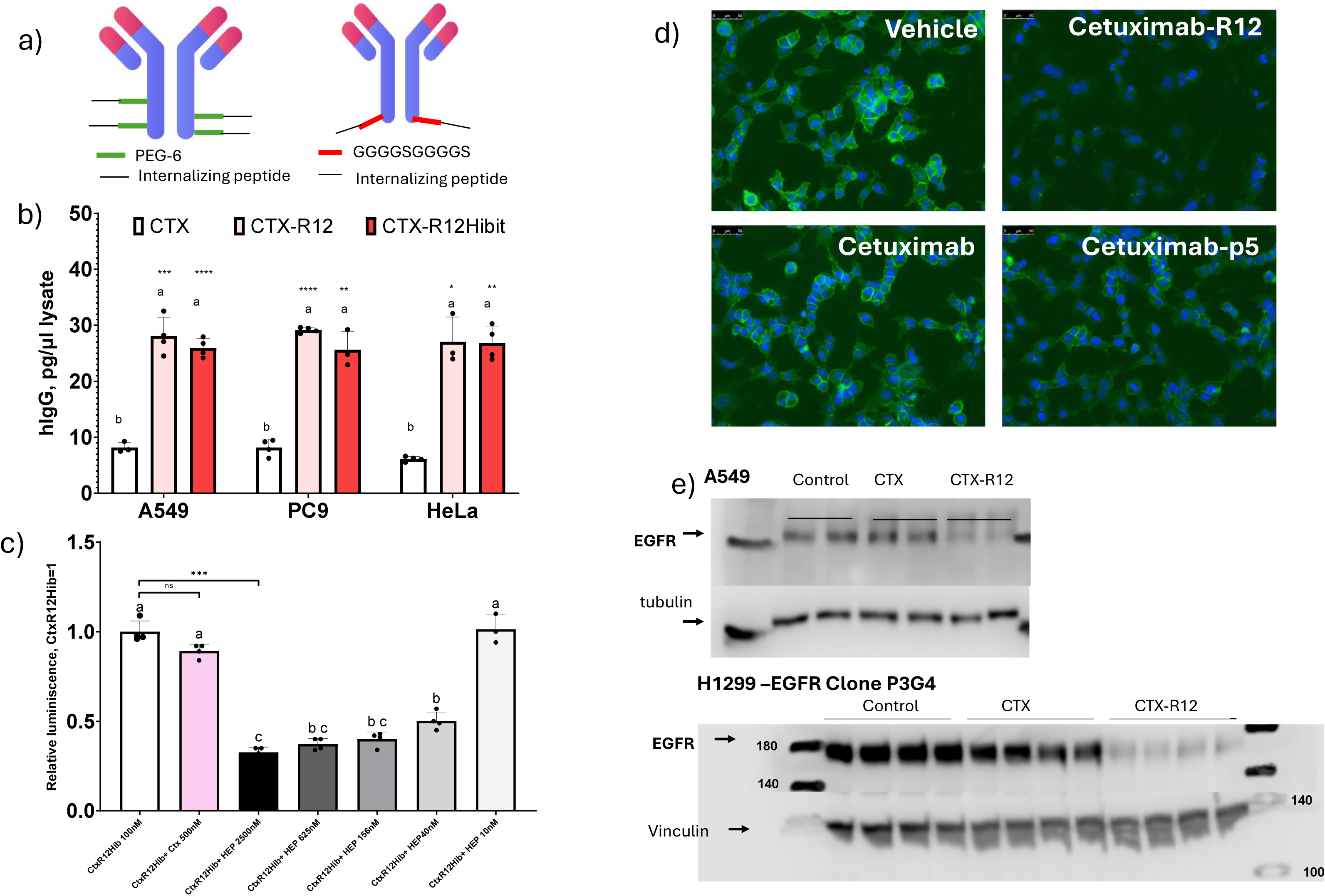
Chemical internalizing-peptide conjugation of CTX enhances CTX internalization and promotes EGFR depletion. **(a)** Schematic of CTX internalizing-peptide chemical conjugation (left) and fusion protein (right) used in this study. **(b)** Intracellular human IgG quantification. A549, PC9 and HeLa cells were treated for 3 h with 100 nM CTX, CTX-R12 or CTX-R12Hibit. Intracellular human IgG was quantified by ELISA and expressed as pg/µL of cell lysate (n = 3-4 per condition). **(c)** Effect of CTX and HEP on CTX-R12Hibit-associated luminescence. A549 cells were treated with CTX-R12Hibit (100 nM) alone or together with CTX (500 nM) or HEP at the indicated concentrations. Luminescence was normalized to CTX-R12Hibit alone (set to 1; n = 4 per condition, except HEP 10 nM, n = 3) b,c: Bars represent mean ± SD; dots indicate individual measurements. Statistical comparisons were performed using Dunnett’s T3 multiple-comparisons test, with all pairwise comparisons included separately within each cell line in (b) and across the seven conditions in (c). Groups sharing at least one letter do not differ significantly; groups with no shared letters differ significantly (adjusted P < 0.05). Letters apply only within each cell line or panel. Asterisks indicate adjusted P values versus CTX in (b) or for bracketed comparisons in (c): *P < 0.05, **P < 0.01, ***P < 0.001 and ****P < 0.0001; ns, not significant. (d) EGFR immunofluorescence in non-permeabilized HeLa cells showing reduced surface-accessible EGFR after CTX-R12 compared with parental CTX and the CTX-p5 control. (e) Total-EGFR immunoblotting of A549 and EGFR-overexpressing H1299 cells following treatment with parental CTX or CTX-R12.

Compared with cells treated for 3 h with parental CTX, cells treated with CTX-R12 and CTX-R12-HiBiT displayed approximately threefold higher intracellular human IgG accumulation (Fig. 1b). In A549 cells, a 25-fold molar excess of soluble heparin markedly reduced intracellular CTX-R12-HiBiT accumulation by approximately 68%, whereas excess unconjugated CTX produced comparatively limited inhibition (Fig. 1c). Notably, heparin did not completely abolish intracellular accumulation, suggesting that a fraction of CTX-R12-HiBiT uptake persists despite competition with soluble heparin. This pattern is consistent with the previously described involvement of heparan sulfate interactions in polyarginine-mediated cellular entry [8] and suggests that R12 conjugation introduces a substantial heparin-sensitive component of antibody uptake beyond the intrinsic endocytic behavior of parental CTX.

To determine whether enhanced CTX-R12 internalization affected EGFR availability in HeLa cells, we examined surface-accessible EGFR by immunofluorescence under non-permeabilizing conditions, using an EGFR-detection antibody that does not compete with CTX for receptor binding (verified by cell-ELISA, supplementary Fig. 9). Treatment with CTX-R12 markedly reduced detectable surface EGFR compared with parental CTX, whereas the conjugate of CTX with the non-internalizing control peptide P5 did not reproduce this effect (Fig. 1d). These observations indicate that surface receptor depletion depends on the properties of the attached peptide and cannot be attributed simply to the chemical modification of the antibody.

Importantly, reduced surface-accessible EGFR was accompanied by decreased total receptor abundance. Whole-cell immunoblotting of A549 cells and EGFR-overexpressing H1299 cells revealed reduced total EGFR levels following CTX-R12 treatment (200nM for 3 days in A549, 100nM in H1299 for 24h), compared with parental CTX (Fig. 1e). Thus, internalizing-peptide conjugate promoted not only altered receptor localization but also measurable loss of total cellular EGFR.

Together, these findings establish CTX-R12 as an antibody-peptide construct capable of enhancing intracellular antibody accumulation and promoting both surface and total EGFR depletion.

### 2. Internalizing-peptide conjugation reduces wound repopulation and requires combined EGFR recognition and internalizing activity

We next investigated whether CTX-R12-mediated EGFR depletion was accompanied by a distinct functional phenotype, as assessed by its effects on wound repopulation. Wound-repopulation assays were performed in serum-starved A549, PC9 and H1975 cells following treatment with EGF (50ng/ml), parental CTX or its internalizing-peptide conjugates for 24/48h.

CTX-R12 consistently produced stronger inhibition of wound closure than parental CTX across the tested cellular models (Fig. 2a), reducing closure toward the levels observed in unstimulated controls. To compare this phenotype with receptor depletion achieved through an independent mechanism, we additionally performed siRNA-mediated EGFR silencing (see Materials and methods). In PC9 and H1975 cells, siEGFR produced a more pronounced inhibition of wound repopulation than CTX-R12 (Fig. 2b). These findings are consistent with previous observations that EGFR knockdown inhibits wound closure and proliferation in H1975 cells [20]. Notably, the intermediate inhibitory phenotype observed with CTX-R12 suggests that internalizing-peptide engineering confers functional effects beyond conventional EGFR blockade, partially resembling those associated with genetic receptor depletion.

**Figure 2.**
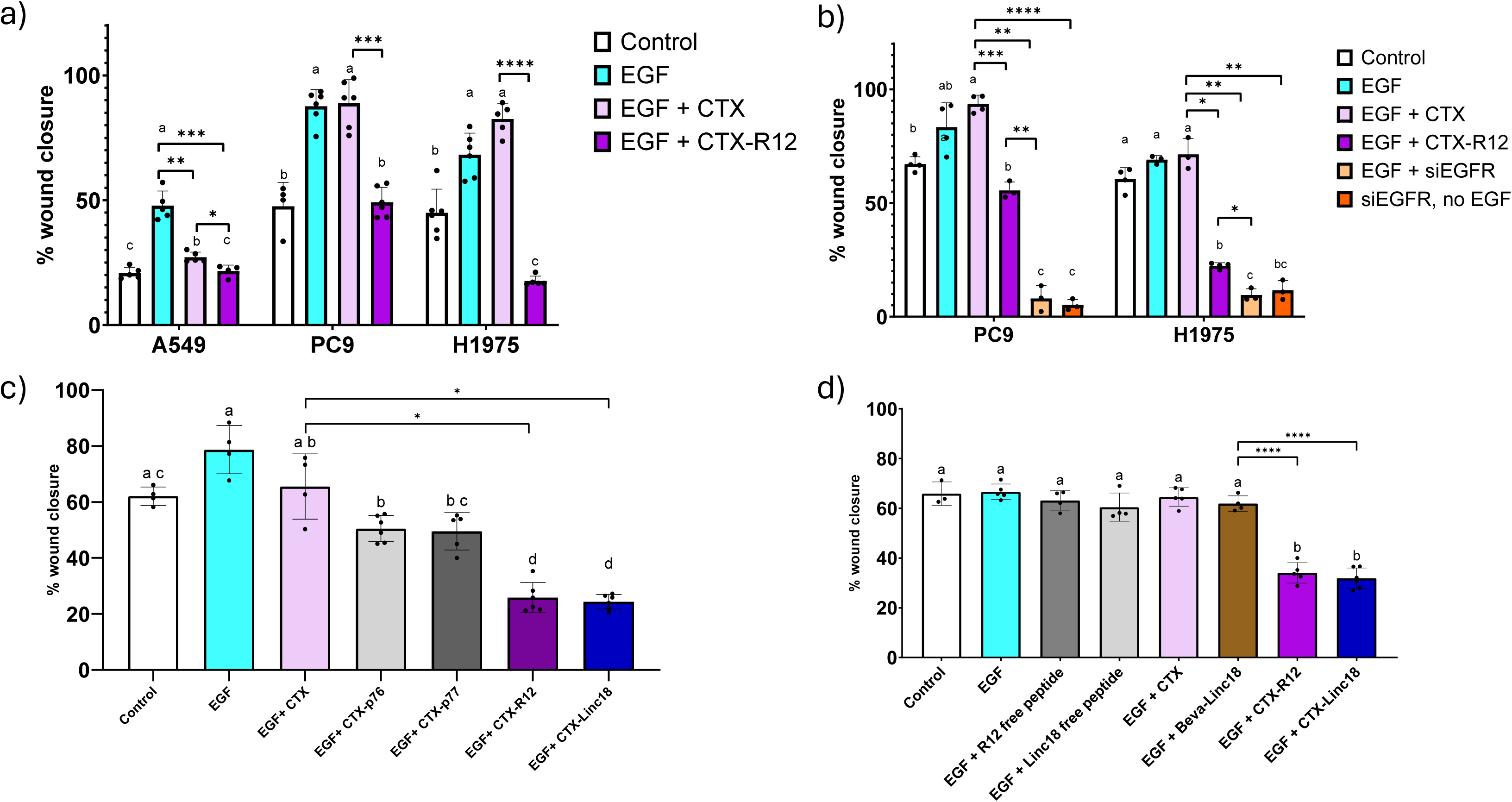
Chemical conjugates of cetuximab with Internalizing peptides enhance inhibition of wound repopulation and require simultaneous presence of EGFR binding moiety and internalizing peptide. (a) Wound-repopulation responses to parental CTX and CTX-R12 in A549, PC9 and H1975 cells. (b) Comparison of CTX-R12 activity with siRNA-mediated EGFR depletion. (c) Functional comparison of CTX-R12, CTX-LINC18 and the poorly internalizing CTX-P76/P77 controls. (d) Control experiments with unconjugated peptides and irrelevant antibody-peptide conjugates. Points represent individual Wells (3-6 replicates) per condition; Bars show mean ± SD. Group means were compared using Welch’s one-way ANOVA followed by Dunnett’s T3 multiple-comparisons test for all pairwise comparisons (family-wise α = 0.05). Letters above groups summarize the adjusted comparisons: groups with no letter in common differ significantly (adjusted P < 0.05), whereas groups sharing at least one letter were not significantly different. *, **, ***, **** Adjusted p-value <0.05, <0.01, <0.001,<0.0001

To determine whether the enhanced activity was specific to R12, we evaluated CTX conjugated to the internalizing peptide LINC18 (a variant of R12), and to the poorly internalizing peptides P76 and P77. CTX-LINC18 and CTX-R12 strongly (by 60%) inhibited wound repopulation in comparison with CTX, whereas CTX-P76 and CTX-P77 did not reproduce the enhanced inhibitory effect (Fig. 2c).

Additional specificity controls included treatment with free R12 and LINC18 peptides and with LINC18 chemically conjugated to the irrelevant anti-VEGF antibody bevacizumab. While in this assay CTX-LINC18 and CTX-R12 reproducibly decreased wound closure, neither the free peptides nor the irrelevant antibody conjugate had any effect (Fig. 2d).

Consistently, non-permeabilized immunofluorescence of H1975 cells used in wound-repopulation experiments demonstrated reduced surface-accessible EGFR following CTX-R12 and CTX-LINC18 treatment, but not after parental CTX treatment (Supplementary Fig. 1).

Collectively, these findings suggest that the enhanced functional phenotype requires the presence of an EGFR-recognition element and an effective internalizing peptide within the same construct, supporting a biological effect beyond conventional receptor blockade.

### 3. Internalizing-peptide engineering reproduces the enhanced functional phenotype in the independent blocking anti-EGFR antibody LEI

To determine whether the effects observed with CTX conjugates could be transferred to another antibody scaffold, we investigated the proprietary full-length human anti-EGFR antibody LEI. The functional blocking activity of parental LEI was assessed using an acute EGF-stimulation experiment. LEI pretreatment abolished EGF-induced EGFR phosphorylation in serum-starved A549 cells while total EGFR remained detectable, demonstrating inhibition of receptor activation (Supplementary Fig. 2, left).

We subsequently evaluated chemically conjugated LEI-R12 and LEI-LINC18 in H1975 wound-repopulation assays. Treatment with 100nM of the both constructs produced stronger inhibition of wound closure than parental LEI, comparable to the inhibition observed with CTX-R12 and CTX-LINC18 (Fig. 3). In contrast, conjugation of the control peptide P56 to LEI did not reproduce the enhanced inhibitory effect (Supplementary Fig. 3).

**Figure 3.**
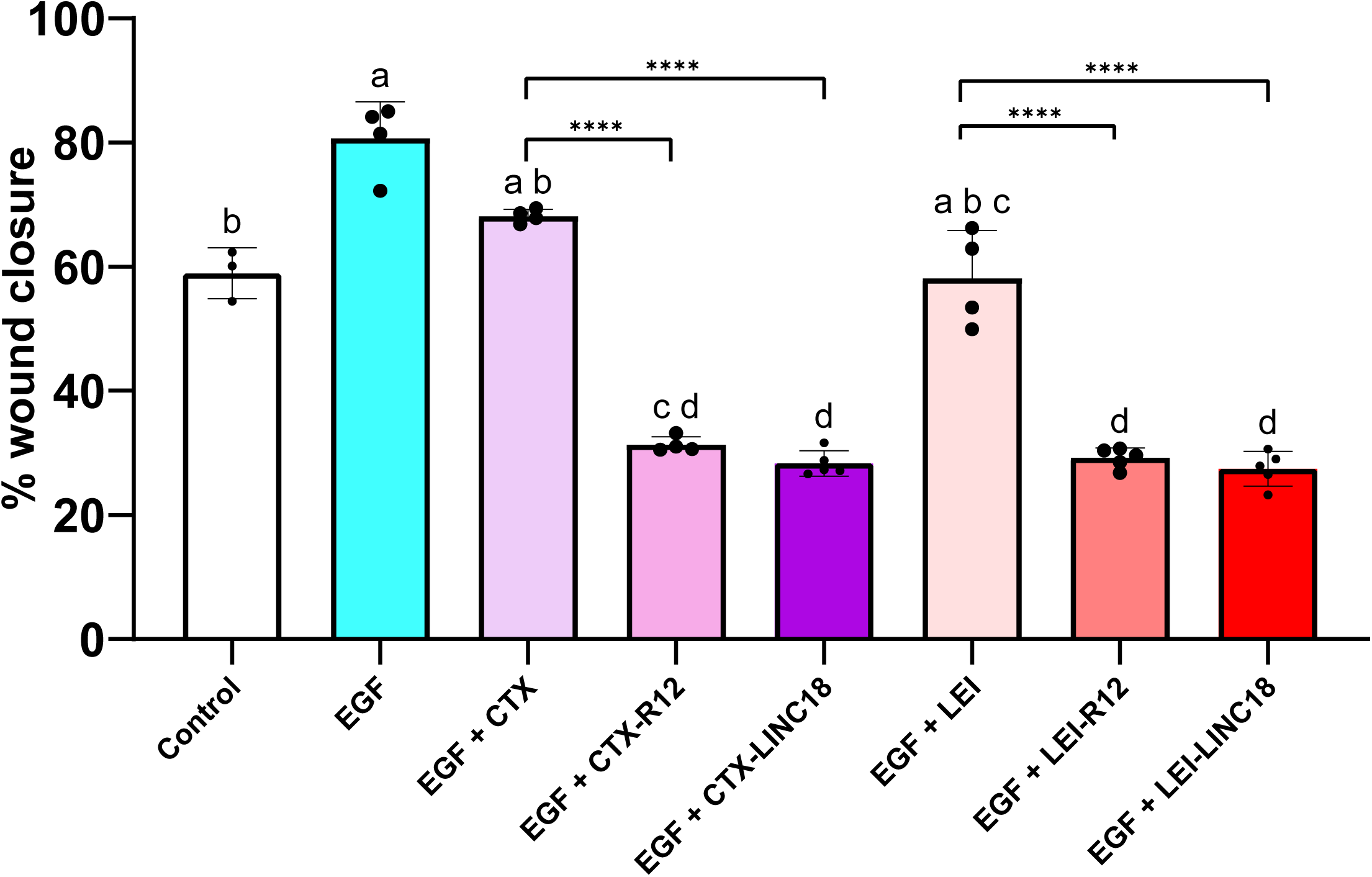
Chemical conjugation of R12 or LINC18 enhances the functional activity of the independent anti-EGFR antibody LEI in a H1975 wound-repopulation experiment comparing parental LEI with chemically conjugated CTX-R12, CTX-LINC18, LEI-R12 and LEI-LINC18. Points represent individual Wells (3-6 replicates) per condition; Bars show mean ± SD. Group means were compared using Welch’s one-way ANOVA followed by Dunnett’s T3 multiple-comparisons test for all 28 pairwise comparisons (family-wise α = 0.05). Letters above groups summarize the adjusted comparisons: groups with no letter in common differ significantly (adjusted P < 0.05), whereas groups sharing at least one letter were not significantly different. **** Adjusted p-value <0.0001

These findings demonstrate that chemical conjugation of selected internalizing peptides enhances the functional activity of a second blocking anti-EGFR antibody, suggesting that the phenotype which the conjugates exert is not restricted only to the CTX scaffold.

### 4. Recombinant anti-EGFR-LINC19 fusion proteins preserve enhanced biological activity

Although chemical conjugation provided an effective method for evaluating internalizing-peptide activity, it requires multiple additional post-expression steps and may generate heterogeneous antibody products. We therefore investigated whether the enhanced activity could be preserved in recombinant full-length antibody fusion proteins.

LINC19, a variant of LINC18, was fused to the C-terminus of the heavy chains of CTX and LEI through a GGGGSGGGGS linker, generating CTX(F)LINC19 and LEI-LINC19, the latter designated LEIP. This design enables the antibody and internalizing peptide to be expressed as a single recombinant molecule without subsequent chemical conjugation (Fig. 1a, right). Similarly to LEI, pretreatment of serum-starved A549 cells with LEIP abolished EGF-induced EGFR phosphorylation (Supplementary Fig. 2, right).

Treatment with 50nM CTX(F)LINC19 for 24h (PC9,H1975) or 48h (A549) retained pronounced inhibition of wound repopulation in PC9, H1975 and A549, cells, comparable to the effects observed with 50nM of chemically conjugated CTX(C)R12 and CTX(C)LINC18 (Fig. 4a; Supplementary Figs. 4-6). Similarly, recombinant LEIP markedly inhibited H1975 wound closure for 24h compared with parental LEI (Fig. 4b), with activity maintained across the tested concentration range of 12.5-100 nM (Fig. 4c).

**Figure 4.**
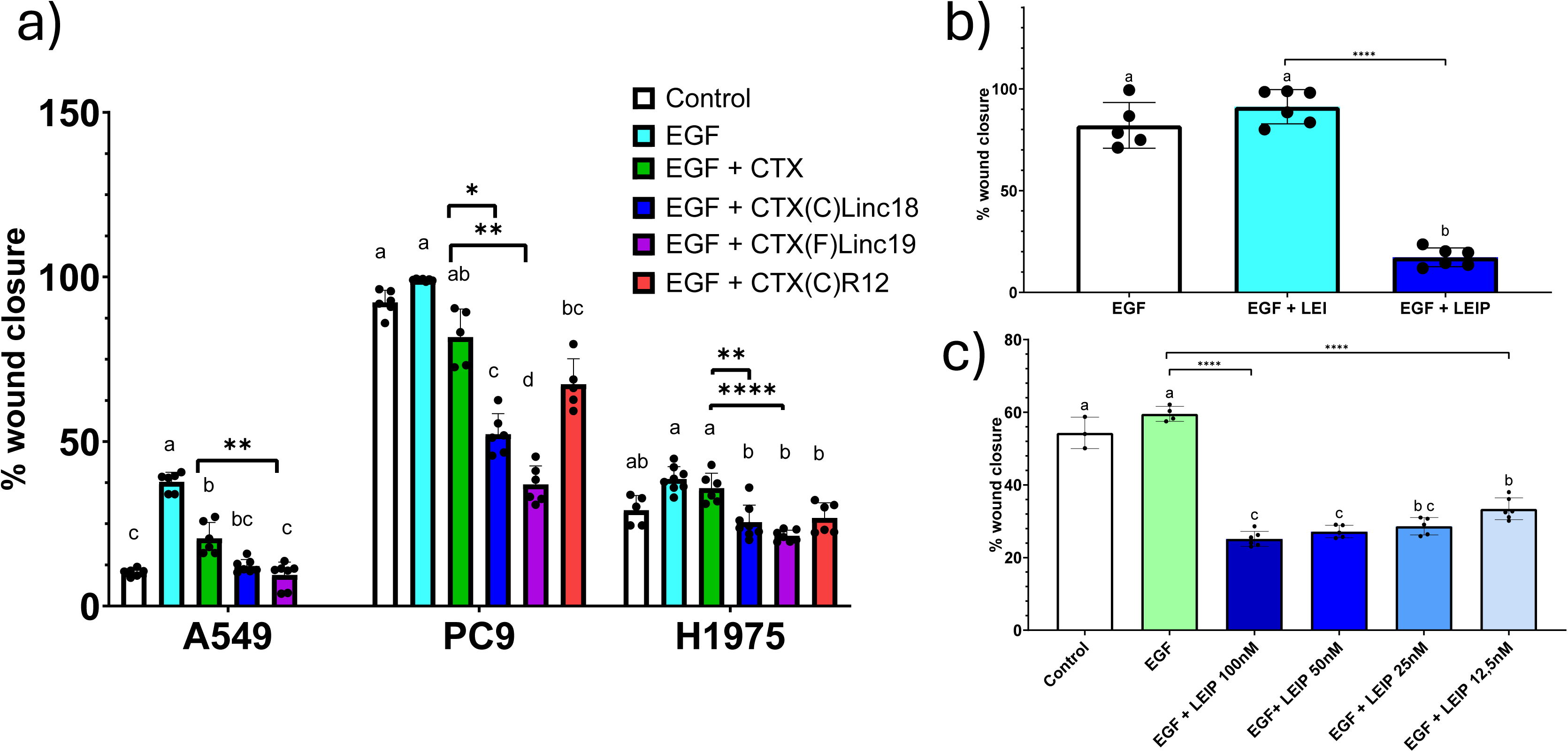
Recombinant anti-EGFR-LINC19 fusion proteins preserve enhanced inhibition of wound repopulation. **(a)** Responses of A549, PC9 and H1975 cells to CTX-based recombinant LINC19 fusion construct; (C) indicates chemical conjugation and (F) indicates fusion. **(b)** Comparison of parental LEI and recombinant LEIP in H1975 wound-repopulation assays. **(c)** Concentration-dependent inhibition of H1975 wound repopulation by recombinant LEIP. Points represent individual wells (3-6 replicates) per condition; Bars show mean ± SD. Group means were compared using Welch’s one-way ANOVA followed by Dunnett’s T3 multiple-comparisons test for all pairwise comparisons (family-wise α = 0.05). Letters above groups summarize the adjusted comparisons: groups with no letter in common differ significantly (adjusted P < 0.05), whereas groups sharing at least one letter were not significantly different. *, **, **** Adjusted p-value <0.05, <0.01, <0.0001

We next investigated whether the enhanced functional phenotype was accompanied by receptor depletion. EGFR immunofluorescence with non-permeabilized A549 and H1975 cells, which were treated for 24h with 100 nM LEI and 100 nM LEIP, demonstrated that LEIP markedly reduced surface-accessible relative to parental LEI (Fig. 5). The reduction in EGFR staining approached that observed following siRNA-mediated EGFR silencing, included as an independent receptor-depletion control.

**Figure 5.**
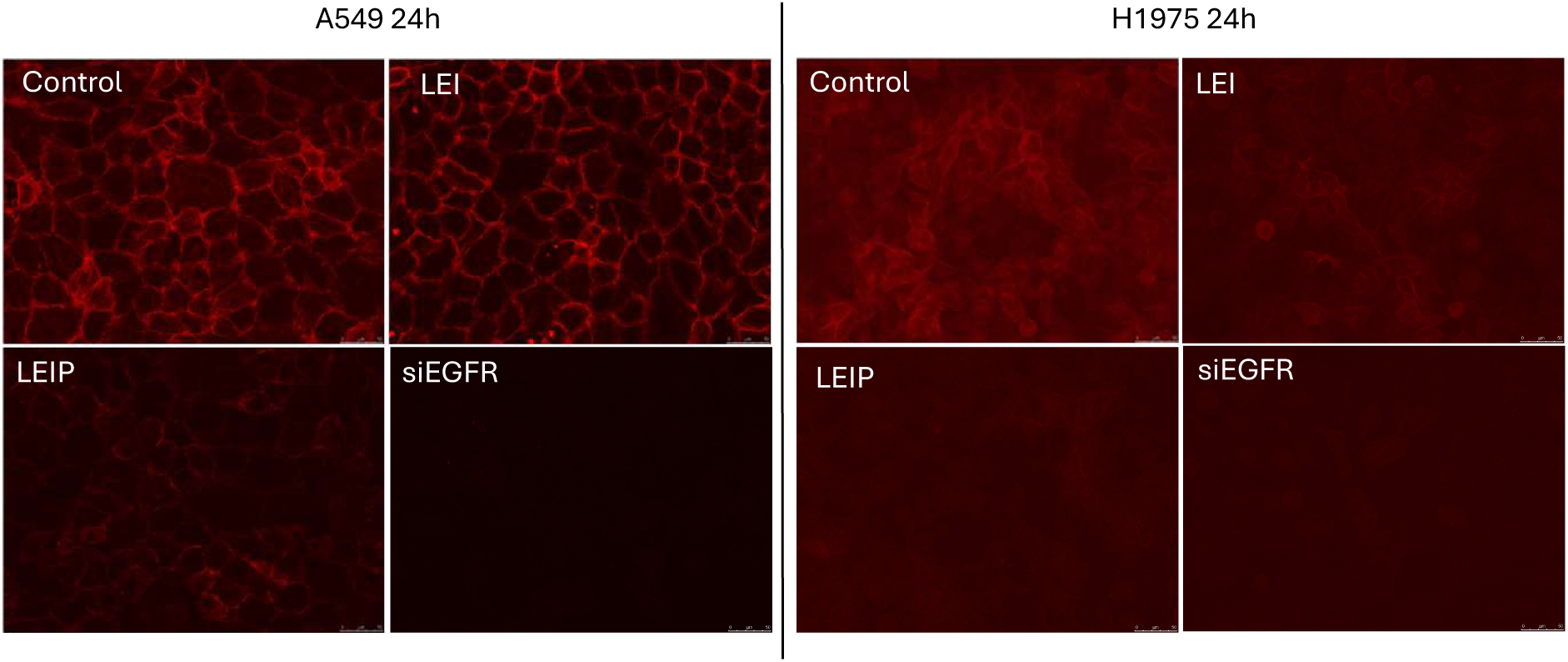
Recombinant LEIP reduces surface-accessible EGFR in A549 and H1975 cells. Representative images of EGFR immunofluorescence in non-permeabilized A549 and H1975 after control, LEI or LEIP treatment. EGFR-silenced cells were included as receptor-depletion controls. Red -mouse anti EGFR 199.12 (detects extracellular domain) of non-permeabilized cells.

Together, these findings establish that the enhanced biological activity associated with internalizing-peptide engineering can be preserved in recombinant full-length antibodies. Moreover, LEIP retains the ability to markedly reduce surface-accessible EGFR, supporting transfer of the receptor-depletion phenotype to a defined recombinant format.

### 5. Internalizing anti-EGFR constructs reduce DNA synthesis, contributing to inhibition of wound repopulation

Because reduced wound closure was repeatedly observed following treatment with internalizing anti-EGFR constructs, we investigated whether this phenotype reflected changes in proliferation, migration or both. Two complementary experimental approaches were employed. Mitomycin C was used to suppress proliferation and determine whether treatment-dependent differences in wound closure persisted under proliferation-restricted conditions, whereas bromodeoxyuridine (BrdU) incorporation was used to independently assess DNA synthesis.

In serum-starved A549 cells, mitomycin C substantially reduced the differences in wound closure between cells treated with the parental CTX and the chemically conjugated CTX(C)LINC18, and CTX(F)LINC19 (Fig. 6a, left versus right), suggesting that cell proliferation contributes to the observed inhibition of wound repopulation. However, this interpretation requires caution, as mitomycin C has been reported to affect wound closure independently of its antiproliferative activity, including in A549 cells [18].

**Figure 6.**
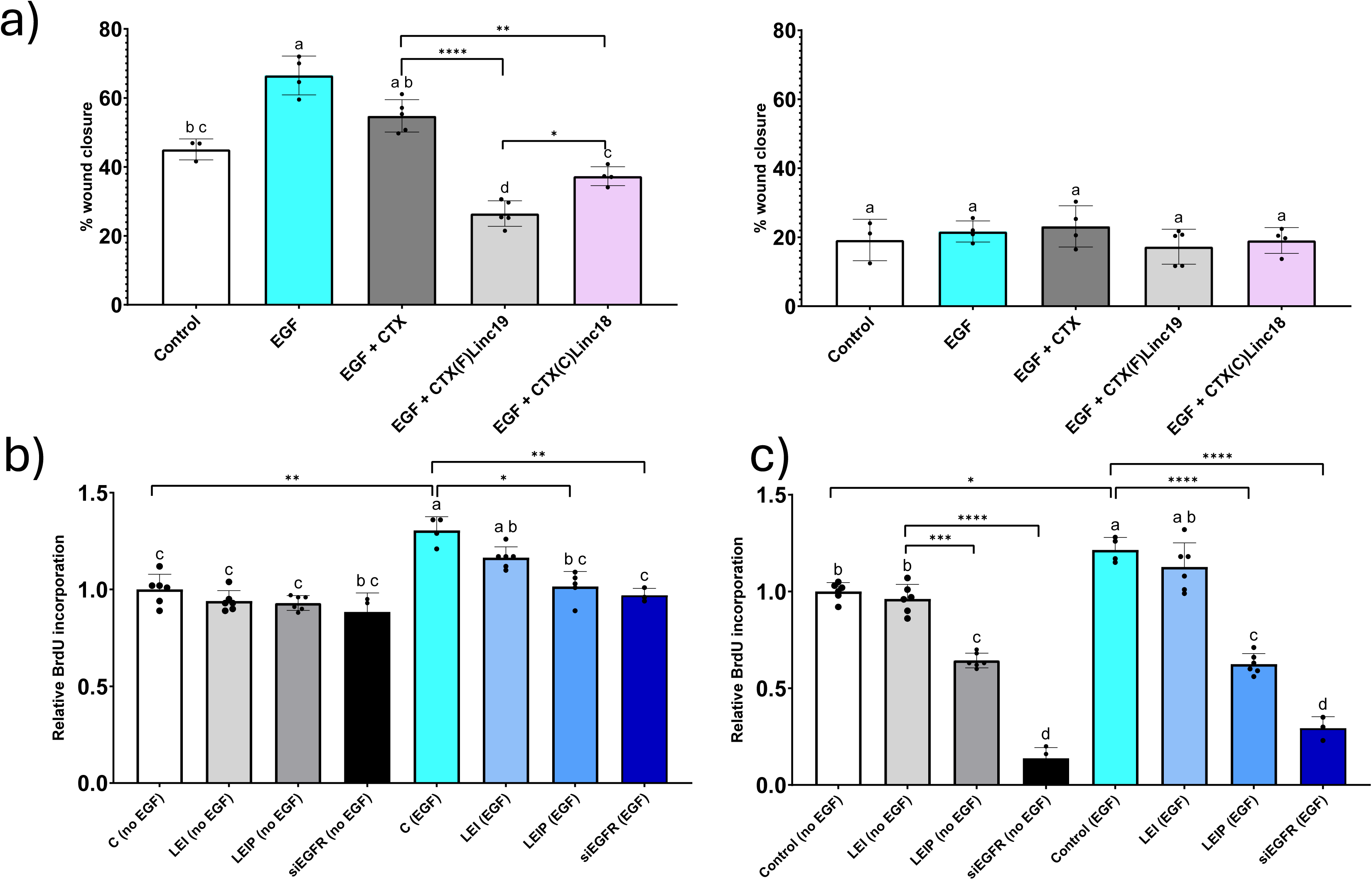
Reduced DNA synthesis contributes to the inhibition of wound repopulation by internalizing anti-EGFR constructs. (a) A549 wound-repopulation experiment without (left) and with mitomycin (right). (b) BrdU incorporation in serum-starved A549 cells following LEI, LEIP and EGFR-silencing treatments ± EGF. (c) BrdU incorporation in serum-starved H1975 cells following LEI, LEIP and EGFR-silencing treatments ± EGF. Points represent individual wells (3-6 replicates) per condition; Bars show mean ± SD. Group means were compared using Welch’s one-way ANOVA followed by Dunnett’s T3 multiple-comparisons test for all pairwise comparisons (family-wise α = 0.05). Letters above groups summarize the adjusted comparisons: groups with no letter in common differ significantly (adjusted P < 0.05), whereas groups sharing at least one letter were not significantly different. *, **, **** Adjusted p-value <0.05, <0.01, <0.0001

Consistent with a contribution from reduced proliferative activity, LEIP treatment produced a modest reduction in BrdU incorporation in serum starved A549 cells stimulated with EGF, similarly to siEGFR silenced cells (Fig. 6b). Similarly, in H1975 cells, both LEIP and siEGFR significantly reduced BrdU incorporation, whereas LEI did not (Fig. 6c). These experiments were performed in serum-starved cells, both in the absence and presence of EGF stimulation, allowing assessment of DNA synthesis under basal and growth-factor-stimulated conditions.

Interestingly, crystal violet assays performed in serum-containing, subconfluent cultures did not reveal a corresponding reduction in adherent cellular biomass following treatment with anti-EGFR constructs over the examined 24-72 h period (Supplementary Fig. 7). This discrepancy may reflect differences in growth-factor availability between the experimental conditions. Under serum-deprived conditions, EGFR depletion may exert a greater influence on DNA synthesis, whereas additional growth factors present in complete medium could sustain cellular growth through alternative signaling pathways. Furthermore, because BrdU incorporation was measured during a 2-h labeling period as a total signal per well, the reduction reflects decreased DNA synthesis during that interval rather than directly demonstrating sustained inhibition of cell proliferation.

Together, these findings support a contribution of reduced DNA synthesis to the inhibitory phenotype associated with internalizing anti-EGFR constructs, while suggesting that the magnitude of their antiproliferative effects may depend on the growth-factor environment. Nevertheless, because mitomycin C can independently influence wound closure, the relative contributions of proliferation and migration cannot be established from these experiments. Accordingly, the term *wound repopulation* is used rather than attributing the observed inhibition exclusively to impaired migration.

### 6. Enhanced internalization of recombinant LEIP is associated with acidification-dependent EGFR degradation

Having established that recombinant LEIP enhances the biological activity of anti-EGFR antibodies and markedly reduces surface-accessible EGFR (Figs. 4-5), we next investigated its intracellular accumulation and whether these effects were accompanied by a reduction in total cellular receptor abundance.

To assess the internalization of recombinant LEIP, A549 cells were treated for 3 h with parental LEI or LEIP labeled with Alexa Fluor 647, a pH-insensitive fluorophore allowing quantification of intracellular antibody-associated fluorescence independently of endosomal acidification. Following conjugation, unreacted fluorophore was removed by centrifugal ultrafiltration using Amicon filters. After treatment, cells were subjected to acidic stripping to remove surface-bound antibodies, followed by cell lysis and fluorescence quantification. Lysates from LEIP-AF647-treated cells exhibited approximately 3.1-fold higher fluorescence than those from LEI-AF647-treated cells (Fig. 7a), closely reproducing the enhanced intracellular accumulation previously observed with chemically conjugated CTX-R12 constructs (Fig. 1b). These findings indicate that the enhanced internalization associated with internalizing-peptide engineering is preserved in the recombinant LEIP format.

**Figure 7.**
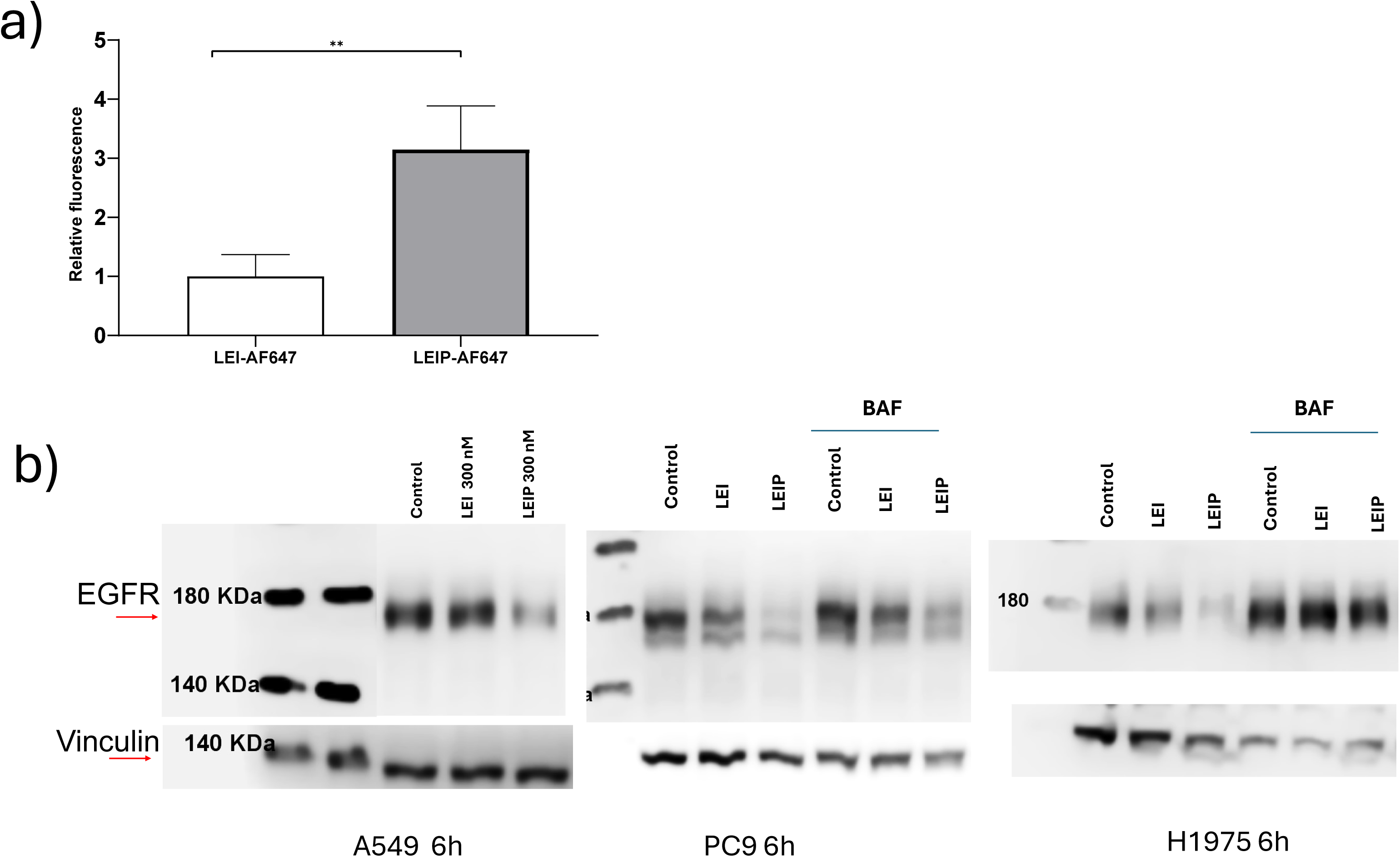
Enhanced intracellular accumulation of recombinant LEIP is associated with bafilomycin-sensitive EGFR depletion. (a) A549 cells were treated for 3 h with parental LEI or recombinant LEIP labeled with Alexa Fluor 647, a pH-insensitive fluorophore. Following treatment, surface-bound antibodies were removed by acidic stripping, and intracellular antibody-associated fluorescence was quantified in cell lysates. Fluorescence signals were normalized to LEI-treated cells. LEIP exhibited approximately 3.1-fold higher intracellular fluorescence than parental LEI. Data are presented as mean ± SD of technical replicates (LEI, n = 3; LEIP, n = 4). Statistical significance was assessed using a two-tailed unpaired t-test with Welch’s correction (t = 5.036, df = 4.578, P = 0.0051; **P < 0.01) (b) Representative Western blots showing total EGFR levels in A549, PC9 and H1975 cells following treatment with parental LEI or recombinant LEIP (100 or 300 nM, as indicated) for 6 h, with or without pretreatment with bafilomycin A1 (BafA1; 100 nM, 1 h). LEIP reduced the abundance of the conventional EGFR band, whereas BafA1 attenuated or reversed this reduction, supporting an acidification-dependent component of EGFR degradation. Corresponding loading controls are shown below the EGFR blots. Molecular weights are indicated in kDa.

We next investigated whether enhanced LEIP internalization was accompanied by EGFR depletion. Whole-cell immunoblotting of A549, PC9 and H1975 cells treated with parental LEI or recombinant LEIP (100/300nM, 6 h) revealed a reduction in the abundance of the conventional EGFR band following LEIP treatment (Fig. 7b). To determine whether receptor loss involved endolysosomal degradation, we examined the effects of bafilomycin A1, an inhibitor of vacuolar H⁺-ATPase-dependent acidification of endosomal and lysosomal compartments [14]. Pretreatment with bafilomycin A1 (100 nM, 1 h) reversed the LEIP-associated reduction in EGFR abundance following 6 h of antibody treatment (Fig. 7b), supporting the involvement of an acidification-dependent endolysosomal degradation pathway.

Collectively, these findings demonstrate that recombinant LEIP retains the enhanced internalization phenotype originally established with chemically conjugated anti-EGFR constructs and promotes a reduction in total EGFR abundance that is sensitive to inhibition of endolysosomal acidification. Together with the reduction in surface-accessible EGFR observed by immunofluorescence (Fig. 5), these observations support a model in which LEIP promotes receptor internalization followed by degradation, rather than merely redistributing EGFR from the plasma membrane into a sustained intracellular pool. The high-affinity interaction between LEIP and EGFR provides a plausible basis for the co-internalization and subsequent lysosomal targeting of antibody-receptor complexes, although their trafficking as an associated complex remains to be directly demonstrated. Furthermore, the variable extent of bafilomycin-mediated rescue suggests that endolysosomal degradation contributes to LEIP-induced EGFR depletion without necessarily accounting for all receptor loss.

### 7. siRNA-mediated EGFR depletion predicts enhanced response to EGFR kinase inhibition

Having established that LEIP promotes EGFR depletion through an acidification-dependent pathway, we next investigated whether reducing receptor abundance could enhance cellular responsiveness to EGFR TKIs.

Because LEIP combines conventional receptor blockade with receptor depletion, we first employed siRNA-mediated EGFR silencing as an independent experimental approach to assess the contribution of EGFR receptor abundance to TKI responsiveness. Previous studies demonstrated that EGFR-mutant NSCLC cells can retain dependence on receptor expression despite resistance to kinase inhibition [9], and that targeted delivery of EGFR-directed siRNA can enhance gefitinib sensitivity [10].

We examined the effects of gefitinib and osimertinib in EGFR-silenced PC9 and H1975 cells, with EGFR knockdown confirmed by fixed-cell ELISA (Supplementary Fig. 8).

Following 48 h of treatment, PC9 cells transfected with siEGFR exhibited significantly lower crystal violet staining than siNON controls under vehicle conditions and at both tested concentrations of gefitinib and osimertinib (Šídák-adjusted P < 0.0001 for all comparisons; Fig. 8, top row). Significant siRNA treatment × drug interactions were detected for gefitinib (P < 0.0001) and osimertinib (P = 0.0002), supporting altered TKI-associated biomass responses following EGFR depletion.

**Figure 8.**
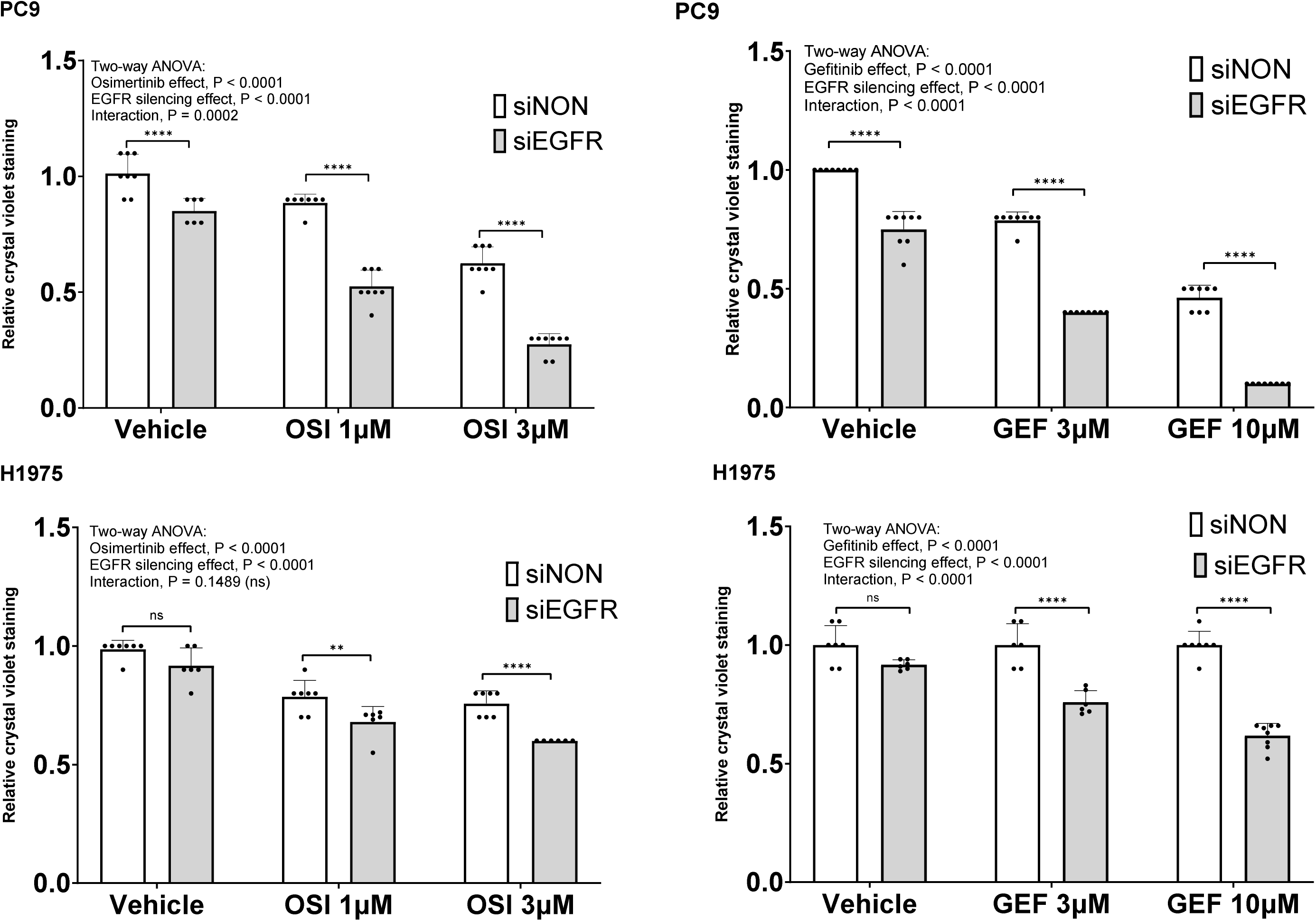
Effects of EGFR silencing on gefitinib and osimertinib responses in PC9 and H1975 cells. Cells transfected with non-targeting siRNA (siNON) or EGFR-targeting siRNA (siEGFR) were treated for 48 h with vehicle, gefitinib (3 or 10 µM), or osimertinib (1 or 3 µM). Adherent cellular biomass was assessed by crystal violet staining and normalized to the corresponding siNON + vehicle control, set to 1. Data are presented as mean ± SD, with 6-8 technical replicate wells per condition; each dot represents an individual well. Statistical analysis was performed separately for each cell line and drug using ordinary two-way ANOVA with silencing and drug treatment as factors, including their interaction, followed by Šídák’s multiple comparisons test. Brackets compare siNON with siEGFR within each treatment condition: ns, not significant; **adjusted P < 0.01; ****adjusted P < 0.0001. ANOVA main-effect and interaction P values are displayed within each plot. Significant silencing × treatment interactions supported enhanced drug-associated biomass reduction for both TKIs in PC9 cells and for gefitinib in H1975 cells. In osimertinib-treated H1975 cells, silencing reduced biomass, but the interaction was not significant (P = 0.1489). EGFR knockdown validation is presented in Supplementary Fig. 8.

In H1975 cells, which harbor the gefitinib resistance-associated EGFR L858R/T790M genotype, gefitinib produced little change in biomass in siNON controls, whereas progressively lower biomass was observed following gefitinib treatment in siEGFR cells. The significant silencing × gefitinib interaction (P < 0.0001), together with lower biomass in siEGFR than siNON cells at both inhibitor concentrations (adjusted P < 0.0001), supports increased gefitinib responsiveness following receptor depletion in this cellular background.

H1975 cells also exhibited lower biomass following EGFR silencing than siNON controls during treatment with osimertinib at 1 µM (adjusted P = 0.0054) and 3 µM (adjusted P < 0.0001). However, the silencing × osimertinib interaction was not statistically significant (P = 0.1489), indicating an additional reduction in biomass without clear evidence of a modified osimertinib response (Fig. 8, bottom row).

Together, these findings support an important contribution of receptor abundance to TKI responsiveness in the tested NSCLC models and provide a genetic rationale for investigating combinations of recombinant LEIP with pharmacological EGFR kinase inhibition.

### 8. Recombinant LEIP enhances the inhibitory effects of EGFR TKIs in H1975 cells

To determine whether LEIP could enhance the inhibitory effects of EGFR TKIs, consistent with the effects observed following siRNA-mediated EGFR depletion, we assessed wound repopulation in serum starved, EGF-stimulated H1975 cells treated with parental LEI or recombinant LEIP, alone or in combination with osimertinib or gefitinib (Fig. 9).

**Figure 9.**
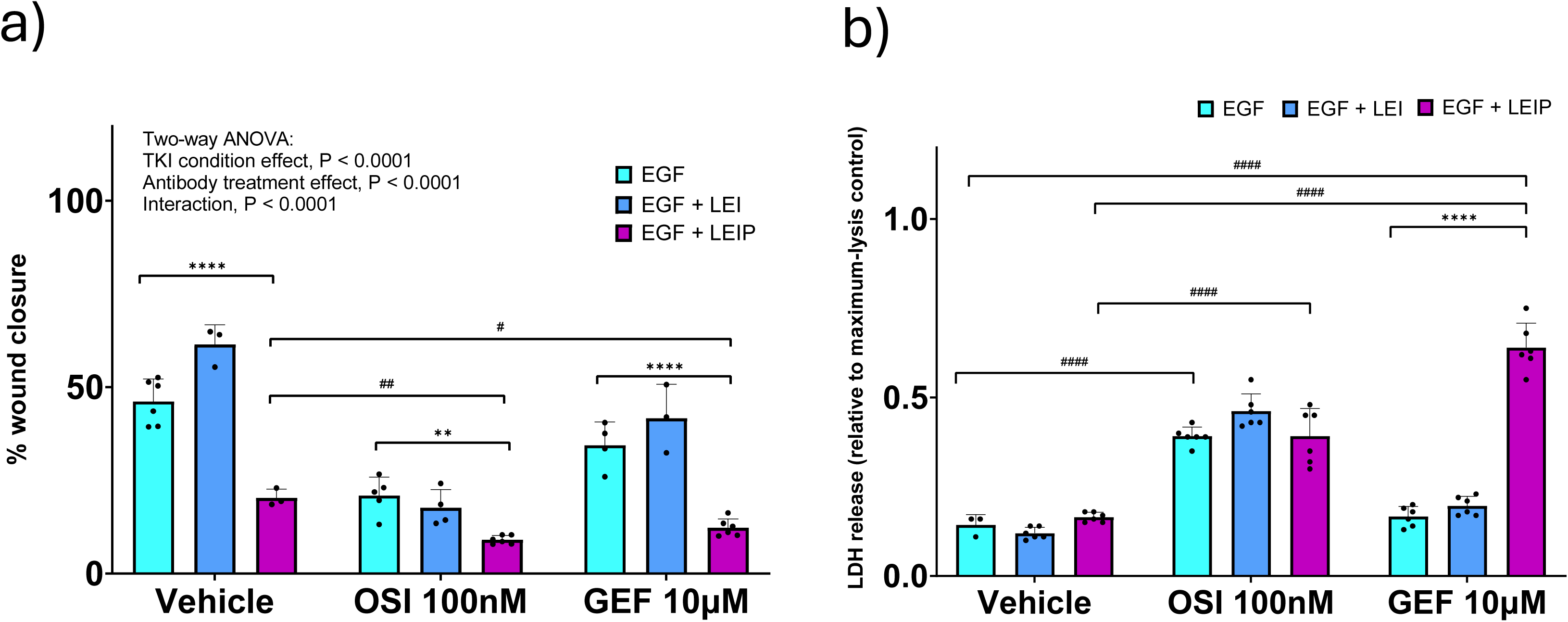
Recombinant LEIP enhances EGFR TKI-associated inhibition of H1975 wound repopulation, with distinct effects on LDH release. (a) Wound closure in EGF-stimulated H1975 cells treated with parental LEI or recombinant LEIP in the absence or presence of osimertinib (OSI, 100 nM) or gefitinib (GEF, 10 µM). (b) LDH activity measured in culture supernatants collected from the same experimental wells at the endpoint of the wound-repopulation assay shown in (a). LDH signals were normalized to the mean signal from three additional wells treated with lysis buffer, assigned a value of 1.0 as the maximum-lysis reference. For each endpoint, statistical analysis was performed using two-way ANOVA followed by Šídák’s multiple-comparisons test for antibody-treatment comparisons within each TKI condition and Dunnett’s multiple-comparisons test for comparisons with vehicle within each antibody-treatment condition. Asterisks indicate Šídák-adjusted significance (**P < 0.01; ****P < 0.0001); hash symbols indicate Dunnett-adjusted significance (#P < 0.05; ##P < 0.01; ####P < 0.0001). Bars represent mean ± SD; individual points represent experimental wells.

In the absence of TKIs, LEIP markedly reduced wound closure compared with both EGF-stimulated controls (20.3% versus 46.2%; Šídák-adjusted P < 0.0001) and parental LEI (20.3% versus 61.5%; adjusted P < 0.0001). Importantly, LEIP also enhanced the inhibitory effects of both TKIs, reducing wound closure compared with osimertinib alone (100 nM; 9.1% versus 20.9%; adjusted P = 0.0033) and gefitinib alone (10 µM; 12.3% versus 34.4%; adjusted P < 0.0001). In contrast, parental LEI did not significantly enhance the effects of either osimertinib or gefitinib (adjusted P = 0.9730 and 0.4264, respectively). Direct comparison of the antibody constructs demonstrated significantly lower wound closure with LEIP than LEI in the presence of gefitinib (12.3% versus 41.7%; adjusted P < 0.0001), whereas their difference in the osimertinib condition was not statistically significant (9.1% versus 17.7%; adjusted P = 0.0915) (Fig. 9a).

To determine whether kinase inhibition provided an additional benefit beyond LEIP treatment alone, we compared wound closure in LEIP-treated cells in the absence and presence of each TKI. Both osimertinib and gefitinib significantly enhanced the inhibitory effect of LEIP, reducing wound closure from 20.3% with LEIP alone to 9.1% and 12.3%, respectively (Dunnett-adjusted P = 0.0050 and P = 0.0475). Two-way ANOVA revealed a significant interaction between antibody treatment and TKI condition (P < 0.0001), indicating that the effects of antibody treatment differed according to the TKI condition (Fig. 9a).

To determine whether the enhanced inhibition of wound repopulation was accompanied by membrane-damage-associated cytotoxicity, culture supernatants were collected from the same experimental wells at the endpoint of the wound-repopulation assay and analyzed for lactate dehydrogenase (LDH) activity (Fig. 9b). LDH signals were normalized to the mean signal from three additional wells treated with lysis buffer, assigned a value of 1.0 as the maximum-lysis reference. In the absence of TKIs, LEIP did not significantly increase relative LDH release compared with EGF-stimulated controls or parental LEI (0.165, 0.143 and 0.120, respectively; Šídák-adjusted P = 0.9975 and P = 0.5318).

Osimertinib alone significantly increased relative LDH release compared with vehicle (0.392 versus 0.143; Dunnett-adjusted P < 0.0001). However, combining osimertinib with LEIP did not produce an additional increase in LDH release compared with osimertinib alone (0.392 versus 0.392; Šídák-adjusted P > 0.9999). Thus, the enhanced inhibition of wound closure observed with LEIP plus osimertinib was not accompanied by an additional increase in membrane-damage-associated LDH release relative to osimertinib alone.

In contrast, gefitinib alone did not significantly increase LDH release compared with vehicle (0.167 versus 0.143; Dunnett-adjusted P = 0.6409), whereas its combination with LEIP resulted in a pronounced increase in relative LDH release (0.640), significantly exceeding both gefitinib alone (0.167) and gefitinib combined with parental LEI (0.197; Šídák-adjusted P < 0.0001 for both comparisons). LDH release was also significantly higher with LEIP plus gefitinib than with LEIP alone (0.640 versus 0.165; Dunnett-adjusted P < 0.0001). This increase was observed in the same wells in which the LEIP-gefitinib combination significantly inhibited wound closure, suggesting that increased membrane damage may contribute to the enhanced inhibitory phenotype. Importantly, because LDH values were normalized to the maximum-lysis reference without correction for spontaneous release, they represent relative LDH activity rather than the percentage of dead cells.

Collectively, these results demonstrate that recombinant LEIP enhances the inhibition of H1975 wound repopulation achieved with either osimertinib or gefitinib, but reveal distinct LDH-release profiles for the two combinations. While LEIP enhanced the effect of osimertinib without an additional increase in LDH release relative to osimertinib alone, its combination with gefitinib produced a pronounced increase in membrane-damage-associated LDH release. These findings suggest that the cellular consequences of combining receptor depletion with kinase inhibition may depend on the TKI employed. Further experiments are required to distinguish the contributions of proliferation inhibition and cell death and to establish whether the observed combined effects represent pharmacological additivity or synergy.

### 9. Internalizing anti-EGFR constructs inhibit NSCLC spheroid growth and enhance responses to EGFR kinase inhibitors

Preliminary experiments in two-dimensional monolayer cultures using crystal violet staining suggested a trend that combining recombinant LEIP with EGFR TKIs produced a greater reduction in adherent cellular biomass than either treatment alone (data not shown). These observations prompted us to investigate whether the enhanced combined activity could also be observed in three-dimensional tumor models. Compared with conventional monolayer cultures, three-dimensional models reproduce important features of solid tumor architecture, including cell-cell interactions, spatial organization and heterogeneous drug accessibility. Moreover, EGFR expression and responsiveness to EGFR-directed treatments can differ substantially between 2D and 3D culture conditions [13]. We therefore evaluated internalizing anti-EGFR constructs during the development and subsequent expansion of PC9 three-dimensional structures and in established H1975 spheroids, with particular emphasis on their activity in combination with gefitinib or osimertinib.

Because PC9 cells formed relatively loose aggregates in ultra-low-attachment plates, we established an alternative three-dimensional model using agarose embedding (see Materials and Methods). CTX-R12 markedly suppressed PC9 colony development, reducing the mean projected colony area by 82.6% relative to untreated controls, compared with a 60.0% reduction following parental CTX treatment (Fig. 10a). Importantly, CTX-R12 produced significantly smaller colonies than CTX alone (Dunnett’s T3-adjusted P < 0.0001), demonstrating enhanced inhibition of three-dimensional colony development following internalizing-peptide conjugation. To determine whether the enhanced inhibitory activity extended to established three-dimensional structures, PC9 colonies were allowed to develop in agarose for 10 days before treatment with parental LEI or recombinant LEIP for 3 consecutive days. Colony growth was evaluated by comparing mean projected colony area at Day 0 and Day 3 in the same experimental wells, with at least 60 colonies analyzed per well. Untreated and LEI-treated colonies exhibited significant increases in mean area (P < 0.0001 for both), whereas LEIP-treated colonies showed only a modest, statistically nonsignificant increase (P = 0.0881). Two-way repeated-measures ANOVA revealed significant effects of time (P < 0.0001), treatment (P = 0.0199) and their interaction (P < 0.0001). At Day 3, LEIP-treated cultures exhibited significantly smaller mean colony areas than both untreated controls (Tukey-adjusted P < 0.0001) and LEI-treated cultures (P = 0.0005) (Fig. 10b). These findings demonstrate that recombinant LEIP substantially restricts the expansion of pre-established PC9 colonies. Together, these results indicate that internalizing anti-EGFR constructs can interfere with both the development and subsequent expansion of PC9 three-dimensional cultures.

**Figure 10.**
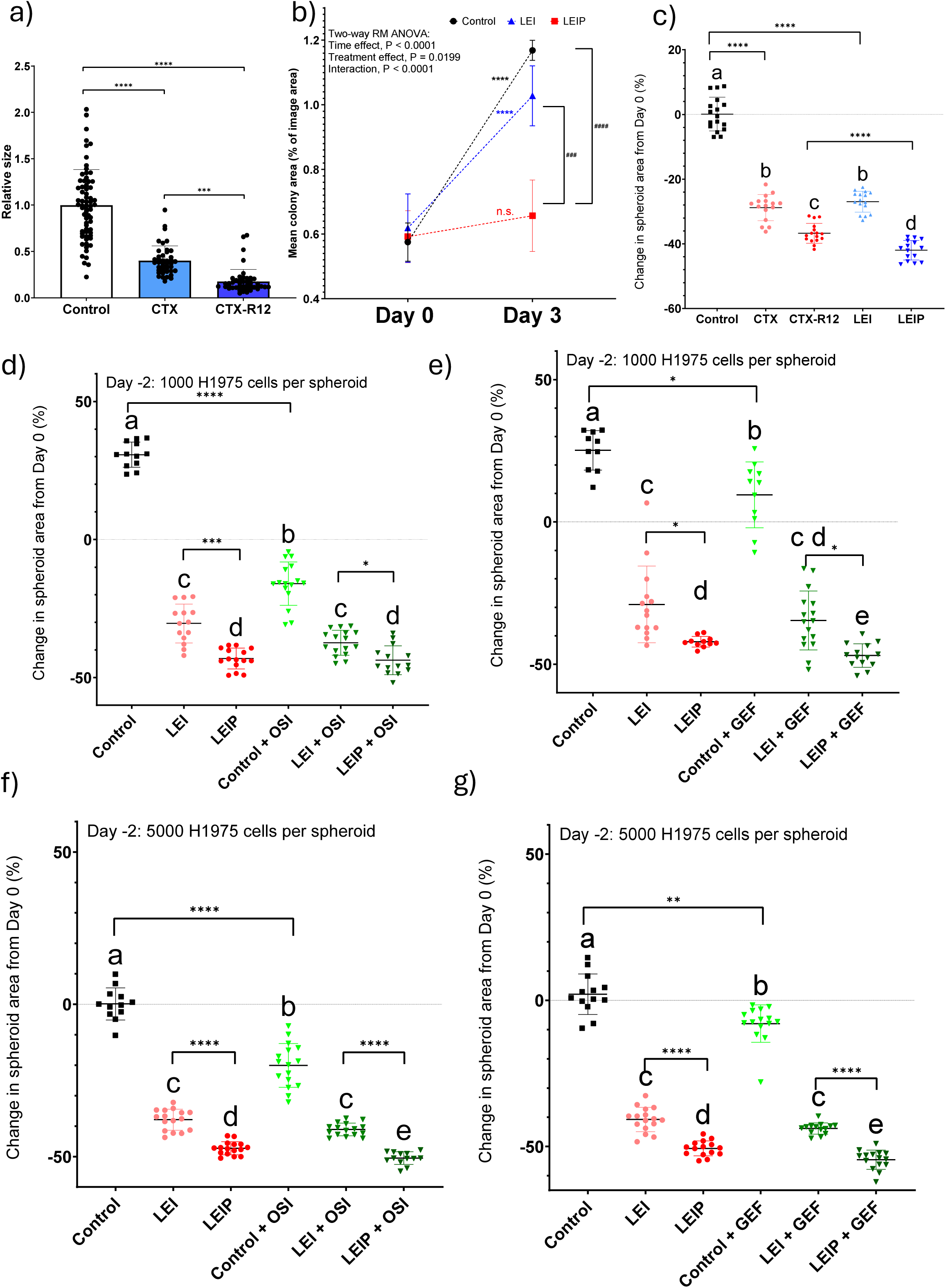
Internalizing anti-EGFR constructs inhibit PC9 colony development and H1975 spheroid growth, retaining activity in combination with EGFR tyrosine kinase inhibitors. **(a)** Effect of parental cetuximab (CTX) and its internalizing-peptide conjugate CTX-R12 on PC9 colony development in agarose. Treatments were applied during colony formation, and projected colony areas were normalized to the mean of the untreated control group (set to 1). Relative mean colony areas were 1.000 for control (n = 68 colonies), 0.400 for CTX (n = 43) and 0.174 for CTX-R12 (n = 48). Data are presented as mean ± SD, with individual colonies shown as points. Statistical analysis was performed using Welch’s one-way ANOVA (W(2, 100.3) = 140.4, P < 0.0001), followed by Dunnett’s T3 multiple-comparisons test. All three pairwise comparisons were significant (adjusted P < 0.0001). Brackets indicate the corresponding comparisons (****P < 0.0001). **(b)** Growth of pre-established PC9 colonies in agarose following treatment with control, parental LEI or recombinant LEIP. Colonies were allowed to develop for 10 days before treatment initiation, and their mean projected area was measured at Day 0 and after 3 days of treatment. At least 60 colonies were analyzed per well, with three replicate wells per treatment condition (n = 3). Data are presented as the mean ± SD of well-level measurements. The same wells were evaluated at both time points, although individual colonies were not tracked longitudinally. Statistical analysis was performed using two-way repeated-measures ANOVA, with time and treatment as factors and measurements matched by well, followed by Tukey’s multiple-comparisons test. Significant effects of time (P < 0.0001), treatment (P = 0.0199) and their interaction (P < 0.0001) were detected. Mean colony area increased significantly between Day 0 and Day 3 in control and LEI-treated cultures (both adjusted P < 0.0001), but not in LEIP-treated cultures (P = 0.0881). At Day 3, LEIP-treated cultures exhibited significantly smaller mean colony areas than untreated controls (P < 0.0001) and LEI-treated cultures (P = 0.0005), whereas control and LEI-treated cultures did not differ significantly (P = 0.1457). Asterisks indicate within-treatment Day 0 versus Day 3 comparisons (****P < 0.0001; ns, not significant), whereas hash symbols indicate between-treatment comparisons at Day 3 (###P < 0.001; ####P < 0.0001). **(c)** Changes in projected area of H1975 spheroids following treatment with parental CTX, CTX-R12, parental LEI or recombinant LEIP. Spheroid area was measured at treatment initiation (Day 0) and after 3 days, and changes were expressed as percentages relative to each spheroid’s initial area. **(d-g)** Effects of parental LEI and recombinant LEIP, alone or in combination with osimertinib (OSI; 0.5 nM) or gefitinib (GEF; 400 nM), on H1975 spheroids initiated with 1,000 (d,e) or 5,000 (f,g) cells per well. Spheroids were allowed to form for 2 days before treatment initiation and were subsequently monitored over 3 days. Panels (d,f) show treatments with or without osimertinib, whereas panels (e,g) show treatments with or without gefitinib. Changes in projected spheroid area were calculated relative to Day 0, with negative values indicating a reduction in projected area. For panels (c-g), individual points represent individual spheroids, and data are presented as mean ± SD. Statistical comparisons were performed using Welch’s one-way ANOVA followed by Dunnett’s T3 multiple-comparisons test. Different lowercase letters indicate statistically significant differences between treatment groups (adjusted P < 0.05), whereas groups sharing at least one letter do not differ significantly. Asterisks indicate the adjusted significance levels of the bracketed comparisons (*P < 0.05; **P < 0.01; ***P < 0.001; ****P < 0.0001). In panels (a) and (b), statistical analyses were performed as described separately above. All percentage changes in spheroid area are relative to the corresponding baseline measurement and do not independently establish changes in viable cellular biomass.

We next investigated the effects of internalizing anti-EGFR constructs in H1975 spheroids generated in ultra-low-attachment plates. The spheroids were allowed to form for 2 days, when they were imaged at Day 0. The typical coefficient of variation in spheroid projected area (µm²) at this stage was less than 5% for all plates used in this work (data not shown), demonstrating excellent uniformity in spheroid size and reproducibility of spheroid formation under the experimental conditions. Treatments were applied during 3 consecutive days and on Day 3, second set of individual images of each spheroid were taken. Initial experiments comparing parental antibodies with their internalizing derivatives demonstrated that CTX-R12 and recombinant LEIP produced greater reductions in projected spheroid area than their respective parental antibodies CTX and LEI (−36.7% for CTX-R12 vs −28.7% for CTX, −41.9% for LEIP vs −27.0% for LEI) (Fig. 10c). These findings extend the enhanced functional activity previously observed in monolayer cultures to a second three-dimensional NSCLC model and support the preservation of this phenotype in the recombinant LEIP format.

To further characterize the response to LEIP and its combination with EGFR TKIs, H1975 spheroids were established using two initial seeding densities, 1,000 or 5,000 cells per well, representing distinct growth states. Control spheroids initiated with 1,000 cells per well exhibited continued expansion (+30.7%), whereas LEI and LEIP markedly reduced their projected area relative to the initial measurement (−30.4 and −43.1%, respectively), LEIP being statistically different than LEI (Fig. 10d). Thus, LEIP not only restricted ongoing spheroid expansion but also produced a measurable decrease in spheroid dimensions under the tested conditions.

Importantly, LEIP-containing combinations with either osimertinib or gefitinib maintained pronounced inhibition of H1975 spheroid growth (Fig. 10d,e). In both experiments, the combinations produced greater reductions in projected area than the corresponding TKI-only conditions and generally stronger responses than combinations containing parental LEI. However, the additional reduction achieved by combining a TKI with LEIP, compared with LEIP alone, was observed only in presence of gefitinib (−42.1 LEIP vs-47.0% LEIP+ GEF, p<0.0146, Fig 10e). Accordingly, these results support the activity of LEIP-containing combinations in growing 3D spheroids without demonstrating that adding a TKI uniformly improves the response beyond LEIP monotherapy.

A distinct growth pattern was observed in larger H1975 spheroids initiated with 5,000 cells per well. Compared with the actively expanding 1,000-cell spheroids, the projected area of the larger, control spheroids, remained essentially the same at day 3 compared with Day 0 (2 days post-seeding). However, LEI and LEIP treatments markedly reduced the projected spheroidś area relative to the initial measurement (−37.9 and −47.3%, respectively), with LEIP being statistically different than LEI (p<0.0001)(Fig. 10f), demonstrating activity in comparatively stable three-dimensional structures. LEIP-containing combinations with osimertinib or gefitinib also produced substantial decreases in spheroid area, exceeding those observed with the corresponding TKI-only and parental LEI combination conditions (LEIP −47.3% vs. LEIP+OSI −50.5%, p<0.006 - Fig. 10f; LEIP −50.7% vs. LEIP+GEF - 54.5%, p<0.02 - Fig. 10g ). These findings indicate that recombinant LEIP retains activity across spheroids with different initial cell densities and growth characteristics. Nevertheless, projected-area measurements cannot independently distinguish loss of viable cellular biomass from structural compaction or disruption of spheroid integrity.

To determine whether pre-existing necrosis could confound the interpretation of treatment-associated changes in spheroid dimensions, and to investigate potential cellular damage following LEIP treatment, H1975 spheroids were stained with Nuclear Green DCS1, a membrane-impermeant fluorescent DNA dye. Control spheroids showed no detectable Nuclear Green-positive material within their cores, indicating the absence of appreciable baseline core necrosis under the experimental conditions (Representative images are shown in Fig. 11a). In contrast, LEIP-treated spheroids exhibited pronounced Nuclear Green fluorescence within the spheroid cores and in material surrounding the spheroid structures. This staining pattern is consistent with treatment-associated loss of membrane integrity and the accumulation of DNA-containing cellular debris. Importantly, these observations indicate that the reduction in spheroid dimensions following LEIP treatment was not confounded by detectable pre-existing core necrosis in the control spheroids. Instead, they suggest that treatment-associated cellular damage may contribute to the observed reduction in projected spheroid area. Nevertheless, the Nuclear Green staining pattern alone does not establish the specific mechanism of cell death or distinguish the relative contributions of cell death and structural changes to spheroid-size reduction.

**Figure 11.**
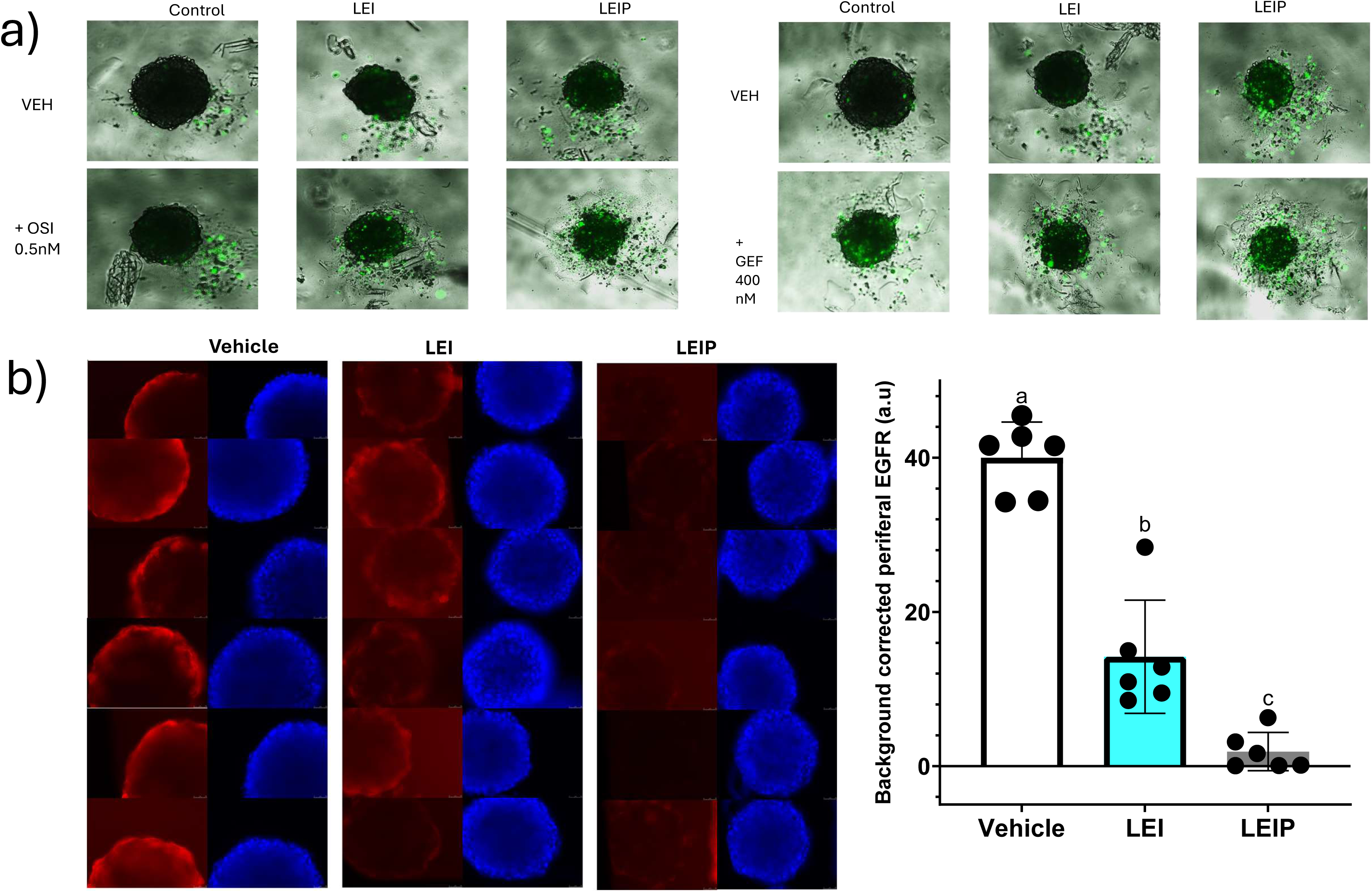
**LEIP treatment is associated with reduced spheroid membrane integrity and diminished peripheral EGFR immunofluorescence. (**a) Representative brightfield/fluorescence overlays of spheroids stained with Nuclear Green DCS1 DNA dye (green), used as a proxy for cell death, following treatment with Control, LEI or LEIP, with vehicle, osimertinib (OSI; 0.5 nM) or gefitinib (GEF; 400 nM). (b) Representative immunofluorescence images of H1975 spheroids showing surface-accessible EGFR (red) and nuclear counterstaining (blue), together with quantification of background-corrected peripheral EGFR fluorescence after treatment with vehicle, LEI, or LEIP. Each point represents one spheroid (n = 6 per group); bars indicate mean ± SD. Statistical differences were assessed using Welch’s ANOVA (P < 0.0001), followed by Dunnett’s T3 multiple-comparisons test. All pairwise comparisons were significant (vehicle vs. LEI, P = 0.0002; vehicle vs. LEIP, P < 0.0001; LEI vs. LEIP, P = 0.0218). Different lowercase letters (a-c) indicate statistically significant differences between groups (adjusted P < 0.05).

Finally, to determine whether the effects of recombinant LEIP in three-dimensional cultures were accompanied by reduced detectable EGFR, H1975 spheroids were treated for 3 days with control, parental LEI or LEIP, fixed, and employed in EGFR and nuclear immunofluorescence analysis. Control spheroids exhibited prominent peripheral EGFR staining, which was reduced following parental LEI treatment and further diminished in LEIP-treated spheroids (Fig. 11b). Quantitative fluorescence analysis confirmed the reduction in detectable EGFR staining following LEIP treatment (Control 40 vs. LEI 14.2 vs LEIP 1.9 a.u), demonstrating that the reduction in detectable EGFR associated with LEIP treatment is also observed at the periphery of three-dimensional H1975 spheroids. Nevertheless, EGFR fluorescence in intact three-dimensional structures can be influenced by antibody penetration, receptor accessibility, spheroid geometry and imaging conditions. Further confocal imaging and quantitative analysis will be required to determine whether the reduction in EGFR staining observed at the spheroid periphery extends throughout the three-dimensional structure.

Collectively, these findings demonstrate that internalizing-peptide-engineered anti-EGFR antibodies inhibit three-dimensional NSCLC outgrowth across different experimental settings, including developing PC9 and pre-established PC9 colonies and H1975 spheroids with distinct growth characteristics (Fig. 10). Recombinant LEIP retained pronounced activity when combined with gefitinib or osimertinib, producing stronger inhibition than corresponding TKI-only or LEI-TKI treatments. The absence of detectable core necrosis in control spheroids, together with treatment-associated Nuclear Green-positive material and reduced EGFR immunofluorescence in LEIP-treated H1975 spheroids (Fig. 11), provides complementary information on cellular integrity and receptor availability in the three-dimensional model. These observations support further investigation of receptor-depleting antibodies as a complementary approach to EGFR kinase inhibition. Formal pharmacological interaction analyses, orthogonal viability measurements and more advanced tumor models will be necessary to establish the therapeutic advantages and mechanistic basis of these combinations.

## Discussion

The present study demonstrates that internalizing-peptide engineering enhances the biological activity of full-length anti-EGFR antibodies by promoting receptor depletion beyond that achieved through conventional antibody blockade. Chemical conjugation of cetuximab (CTX) to R12 increased intracellular antibody accumulation, reduced surface-accessible and total EGFR abundance, and enhanced inhibition of wound repopulation. These effects were reproduced with LINC18 and transferred to the independent anti-EGFR antibody LEI. Importantly, recombinant incorporation of LINC19 preserved the enhanced internalization and functional phenotype, with the resulting LEI-LINC19 fusion (LEIP) promoting acidification-dependent EGFR depletion. The inhibitory activity of these constructs extended to three-dimensional NSCLC models, including PC9 cells harboring an EGFR exon 19 deletion and H1975 cells expressing the L858R/T790M double-mutant receptor. Together with the enhanced responses observed when LEIP was combined with EGFR TKIs, these findings suggest that internalizing-peptide-mediated receptor depletion may provide a complementary approach for targeting EGFR-mutant NSCLC, including cellular contexts characterized by resistance to conventional kinase inhibition.

The distinction between inhibition of EGFR activity and depletion of the receptor protein is central to this approach. Although cetuximab-related antibody-EGFR complexes can undergo substantial endosomal recycling [2], antibody combinations recognizing noncompetitive EGFR epitopes have been shown to promote receptor downregulation by interfering with recycling [3]. Furthermore, cetuximab itself has previously been reported to downregulate mutant EGFR and inhibit the growth of H1975 cells expressing EGFR L858R/T790M [23]. More recently, targeted degradation platforms such as LYTACs and KineTACs have demonstrated that extracellular and membrane-associated proteins can be redirected toward degradation through the recruitment of endogenous trafficking receptors [6,7], with related approaches targeting drug-resistant membrane proteins [15,16]. In contrast, our strategy incorporates an internalization-driving peptide directly into the EGFR-binding antibody without deliberately recruiting a second trafficking receptor. The enhanced EGFR depletion observed with CTX-R12 and LEIP relative to their respective parental antibodies, together with the failure of poorly internalizing peptides, free peptides and an irrelevant antibody conjugate to reproduce the functional phenotype, supports the importance of combining EGFR recognition and internalizing activity within the same molecule.

An important advance of this study is the preservation of enhanced internalization and receptor-depleting activity in the recombinant LINC19 fusion format. Unlike chemical conjugation, genetic incorporation of the internalizing peptide provides predefined peptide positioning and stoichiometry while eliminating additional post-expression conjugation steps. Consistent with previous work demonstrating the feasibility of full-length antibody-cell-penetrating-peptide fusions [5], recombinant LEIP retained enhanced inhibition of wound repopulation and markedly reduced surface-accessible EGFR. Moreover, Alexa Fluor 647-based measurements revealed approximately threefold higher intracellular accumulation of LEIP than parental LEI, closely reproducing the phenotype observed with CTX-R12. Importantly, LEIP-associated loss of total EGFR was attenuated or reversed by bafilomycin A1, supporting the involvement of acidification-dependent endolysosomal degradation [14]. These findings suggest that internalizing-peptide engineering alters the intracellular fate of antibody-associated EGFR, potentially favoring receptor degradation over recycling. Nevertheless, direct demonstration of LEIP-EGFR co-trafficking, the precise sorting mechanisms involved and the kinetics of receptor degradation remain necessary to establish this model.

The functional consequences of internalizing-peptide-mediated EGFR depletion appear to involve alterations in cellular proliferation and wound repopulation. BrdU incorporation experiments demonstrated reduced DNA synthesis following treatment with selected internalizing constructs under serum-starved conditions, while mitomycin C diminished treatment-dependent differences in wound closure. These findings suggest that reduced proliferative activity contributes to the inhibitory phenotype, although the potential effects of mitomycin C on cell motility prevent a definitive separation of proliferation and migration [18]. Notably, crystal violet assays performed in serum-containing cultures did not reveal corresponding reductions in adherent cellular biomass during the examined treatment periods. This discrepancy suggests that the functional consequences of EGFR depletion depend, at least in part, on the growth-factor environment and the experimental endpoint. Accordingly, the reduction in wound closure is best interpreted as inhibition of wound repopulation rather than evidence of an exclusively antimigratory effect.

The distinct EGFR mutational backgrounds of PC9 and H1975 cells provide an important context for interpreting these findings. PC9 cells harbor an activating EGFR exon 19 deletion (E746_A750del), which confers high sensitivity to EGFR TKIs, whereas H1975 cells express the activating L858R mutation together with T790M, a secondary mutation associated with resistance to first-generation inhibitors such as gefitinib [22]. Mechanistically, T790M increases the ATP-binding affinity of mutant EGFR, reducing the effectiveness of reversible ATP-competitive inhibitors [21]. In contrast, strategies that reduce receptor abundance may potentially circumvent this resistance mechanism by eliminating the target protein rather than relying exclusively on inhibition of its catalytic activity. Consistent with previous studies demonstrating that EGFR-mutant NSCLC cells can remain dependent on receptor expression despite resistance to kinase inhibition [9,10], siRNA-mediated EGFR depletion enhanced TKI-associated reductions in cellular biomass in PC9 cells and significantly modified gefitinib responsiveness in H1975 cells. Notably, the interaction between EGFR silencing and gefitinib was significant in H1975 cells, whereas the corresponding interaction with osimertinib was not. These observations suggest that reducing EGFR abundance may partially counteract the functional consequences of T790M-associated gefitinib resistance, although they do not establish complete restoration of drug sensitivity.

The enhanced activity of recombinant LEIP in H1975 cells provides further support for the potential of receptor depletion to complement kinase inhibition in a T790M-positive background. Although H1975 cells are relatively resistant to gefitinib, combining LEIP with this inhibitor reduced wound closure substantially compared with gefitinib alone, whereas parental LEI did not significantly improve the response. Importantly, the LEIP-gefitinib combination also produced greater inhibition than LEIP monotherapy, suggesting that receptor depletion and kinase inhibition exert complementary effects under the experimental conditions. These findings are particularly relevant because LEIP targets extracellular EGFR rather than the ATP-binding pocket affected by the T790M resistance mutation. The enhanced inhibition of wound repopulation was accompanied by a pronounced increase in LDH release with LEIP-gefitinib, suggesting that treatment-associated membrane damage may contribute to the combined phenotype. In contrast, combining LEIP with osimertinib further reduced wound repopulation without increasing LDH release relative to osimertinib alone. Because parental H1975 cells retain sensitivity to osimertinib, this latter observation represents enhancement of an active treatment rather than reversal of established osimertinib resistance. Collectively, these findings suggest that internalizing-peptide-engineered anti-EGFR antibodies may partially counteract the gefitinib-resistant phenotype associated with EGFR T790M, although formal demonstration of pharmacological resensitization will require dose-response analyses and additional viability endpoints.

The activity of internalizing anti-EGFR constructs in three-dimensional NSCLC models further supports the biological relevance of this approach. In PC9 cells, which harbor a TKI-sensitizing EGFR exon 19 deletion, CTX-R12 markedly inhibited the development of agarose-embedded colonies, whereas recombinant LEIP significantly restricted the subsequent expansion of established colonies compared with parental LEI and untreated controls. These observations suggest that receptor-depleting antibodies can interfere with both the formation and continued growth of EGFR-dependent three-dimensional tumor structures. More importantly in the context of kinase-inhibitor resistance, recombinant LEIP also exhibited pronounced activity against H1975 spheroids expressing EGFR L858R/T790M. LEIP reduced the projected area of spheroids initiated with either 1,000 or 5,000 cells per well, producing greater reductions than parental LEI despite their distinct growth characteristics. LEIP-containing combinations with gefitinib or osimertinib produced stronger responses than the corresponding TKI-only treatments. The additional benefit of incorporating a TKI relative to LEIP monotherapy was statistically significant as additional reductions in spheroid area were observed for the gefitinib + LEIP combination in growing and in established spheroids, and osimertinib + LEIP combination in established spheroids. Thus, these findings support substantial independent activity of LEIP in a T790M-positive three-dimensional model, while suggesting that the advantages of combining receptor depletion with kinase inhibition may depend on spheroid growth state and the inhibitor employed. Formal pharmacological interaction analyses remain necessary to distinguish complementary activity from additivity or synergy [17].

Additional observations from H1975 spheroids suggest that the reduction in projected area following LEIP treatment may involve changes in cellular integrity beyond inhibition of spheroid expansion. Nuclear Green DCS1 staining revealed no detectable necrotic cores in untreated spheroids, reducing the likelihood that pre-existing central necrosis confounded the interpretation of treatment-associated changes in spheroid dimensions. In contrast, LEIP-treated spheroids exhibited pronounced DNA-associated fluorescence within and surrounding the spheroid structures, consistent with treatment-associated membrane damage and the accumulation of cellular debris (Fig. 11a). Furthermore, quantitative immunofluorescence demonstrated markedly reduced detectable EGFR in LEIP-treated spheroids compared with parental LEI and untreated controls, supporting persistence of the receptor-depletion phenotype in three-dimensional cultures (Fig. 11b). Nevertheless, spheroid-area measurements and fluorescence imaging cannot independently establish the extent of viable cellular biomass loss or total receptor degradation throughout the spheroid.

Future studies should therefore combine quantitative viability and receptor-degradation measurements with pharmacological sensitivity analyses, including dose-response experiments in H1975 cells and independently established gefitinib- or osimertinib-resistant NSCLC lines. Direct assessment of antibody-EGFR co-trafficking, signaling inhibition, and the selectivity and stability of the recombinant constructs will also be important. Ultimately, in vivo safety and efficacy studies will be required to establish whether internalizing-peptide-mediated EGFR depletion can translate into therapeutic benefit. Nevertheless, the activity demonstrated in both TKI-sensitive and T790M-positive NSCLC models provides a rationale for further investigating receptor-depleting antibodies as a strategy to complement kinase inhibition and potentially overcome selected forms of EGFR-TKI resistance.

## Materials and Methods

### Cell culture

A549 and HeLa cells were maintained in high-glucose DMEM supplemented with 10% fetal bovine serum (FBS), whereas PC9, H1975 and H1299 NSCLC cells were maintained in RPMI 1640 supplemented with 10% FBS. Cultures were maintained at 37 °C in a humidified atmosphere containing 5% CO₂. Where indicated, cells were serum-starved in basal medium supplemented with 0.1% bovine serum albumin (BSA) before stimulation with recombinant EGF. Periodic mycoplasma testing was performed using the Mycoplasma Gel Form kit (90.021-4542; Biotools, Spain).

### Antibodies and internalizing-peptide constructs

Cetuximab (CTX) was used as the reference anti-EGFR monoclonal antibody. LEI is a proprietary full-length anti-EGFR monoclonal antibody developed by Leitat Technological Center using phage display technology. All peptides were synthesized by GenScript. The internalizing peptides R12 and LINC18 were and chemically conjugated to the indicated antibody scaffolds. Control constructs included antibodies conjugated to the peptides P5, P56, P76 or P77, unconjugated peptides and LINC18 conjugated to the irrelevant anti-VEGF antibody bevacizumab.

### Chemical conjugation of antibodies and peptides

The internalizing peptides were chemically conjugated to the indicated antibodies using a maleimide-PEG₆-NHS ester linker. Antibodies were reacted with the NHS ester group of the linker through lysine residues. The linker was dissolved in DMF at 50 mg/mL. One molar equivalent of antibody was mixed with four molar equivalents of linker and incubated for 50 min at 25 °C with shaking at 500 rpm in a thermomixer. The linker solution constituted 5% of the total reaction volume, and the reaction was performed at pH 7. Excess unreacted linker was removed by centrifugal ultrafiltration and buffer exchange using an Amicon Ultra-0.5 centrifugal filter unit with a 10-kDa molecular-weight cutoff. The buffer was exchanged to pH 7.5. The peptide was subsequently conjugated to the maleimide group through its free cysteine residue. Peptides were dissolved in phosphate buffer at 2 mg/mL. One molar equivalent of the antibody-PEG₆-maleimide intermediate was mixed with four molar equivalents of peptide and incubated overnight at 25 °C with shaking at 500 rpm. Excess unreacted peptide was removed by centrifugal ultrafiltration and buffer exchange.

### Acid-stripping fluorescence internalization assay

The internalization capacity of a panel of peptides discussed in this work were preliminarily evaluated in a model assay by conjugating them to anti VEGF antibody bevacizumab (BEVA) and treating A549 cells with fluorescently labelled VEGF alone or in combination with BEVA-peptide conjugates. At the end of the assay, green CellTracker-derived fluorescent normalization reagent (gCDFA) was added for cell normalization. In short, the cellular uptake of VEGF AlexaFluor647 was quantified by acid stripping of the surface-bound material, lysing the cells, and normalizing intracellular fluorescence to a cell-number-dependent green fluorescent signal. For a peptide to be classified as “internalizing”, we set a cut off of at least fivefold high intracellular VEGF fluorescence in cells treated with BEVA-peptide conjugates compared with the negligible baseline internalization of VEGF alone and with BEVA-driven VEGF (data not shown).

### Recombinant expression of antibody fusion proteins

LINC19, a variant of LINC18, was fused to the C-terminus of the antibody heavy chains through a GGGGSGGGGS linker.The resulting recombinant constructs were CTX(F)LINC19 and LEI-LINC19 (LEIP). Recombinant antibody production in CHO cells and purification was performed by GenScript.

### Antibody internalization

Cells were treated with the indicated antibody constructs in the presence or absence of soluble heparin or competing parental antibody. Intracellular antibody accumulation was quantified using the corresponding experimental readout following glycine-based acidic stripping to remove extracellularly associated antibody in two consecutive acid-stripping steps using 100 µL ice-cold 1 M glycine/150 mM NaCl, pH 2.5, for 2 min, followed by neutralization with 10 µL Tris, pH 8.0, and PBS washing after each strip, followed by lysis in RIPA buffer 1x. Intracellular human IgG was quantified by Human IgG (Total) Uncoated ELISA Kit ( 88-50550-88, ThermoFisher Scientific). Intracellular Hibit tag was quantified by Nano-Glo® HiBiT Lytic Detection System (N3030, Promega).

### Surface EGFR immunofluorescence

Cells were fixed with 4% paraformaldehyde for 5 min, blocked with 3% BSA in PBS and stained without membrane permeabilization to assess surface-accessible EGFR.

The EGFR-detection antibody was mouse clone 199.12 (MS-396-P1; Epredia), recognizing an EGFR epitope that does not compete with CTX binding (previous data and our own validation, see Supplementary Figure 9). An Alexa Fluor-labeled anti-mouse secondary antibody was used for detection.

### Whole-cell EGFR immunoblotting

Following treatment, cells were detached enzymatically with trypsin and washed with PBS. Whole-cell lysates were prepared using RIPA lysis buffer supplemented with protease and phosphatase inhibitors.

Protein concentrations were quantified using a BCA assay (23227, Thermo Fisher Scientific). Three volumes of sample were mixed with one volume of 4× Laemmli sample buffer supplemented with β-mercaptoethanol and heated at 70 °C for 10 min.

Equal amounts of protein (10-20 µg) were separated by SDS-PAGE using 6% acrylamide gels. EGFR was detected using EGFR antibody A-10 (sc-373746; Santa Cruz Biotechnology). Phospho EGFR ( Tyr1173) was detected using EGFR antibody 9H2 ( sc-57545; Santa Cruz Biotechnology) Appropriate loading controls were included for comparison between treatment conditions (vinculin antibody (7F9): sc-73614; alpha Tubulin antibody (B-7): sc-5286, Santa Cruz Biotechnology).

### Bafilomycin A1 experiments

To investigate the contribution of endolysosomal acidification to LEIP-associated EGFR loss, cells were pretreated with bafilomycin A1 (100 nM) for 1 h, followed by the addition of LEI or LEIP (100 nM) for a further 6 h.

### EGFR silencing and fixed-cell ELISA

Cells were reverse-transfected with EGFR-targeting siRNA (Silencer Select, catalog 4392420, siRNA ID s564; Thermo Fisher Scientific). All not-siEGFR treated cells were transfected in parallel with a non-targeting control siRNA (Silencer Select Negative Control No. 2), using Lipofectamine RNAiMAX (13778075; Thermo Fisher Scientific).

EGFR knockdown was validated by fixed-cell ELISA before downstream functional analyses (Supplementary Fig. 8). Cells were fixed with 4% paraformaldehyde for 5 min, blocked with 3% BSA in PBS for 1 h and incubated with cetuximab in 3% BSA-PBS containing 0.05% Tween for 1 h at room temperature. Following washing, cells were incubated with an anti-human HRP-conjugated antibody for 1 h, followed by TMB soluble substrate for detection. Plates were read, and cells were further washed and stained with crystal violet for ELISA normalization.

### Wound-repopulation assays

Cells were seeded in full media to reach approximately 90-95% confluence on the next day, before serum deprivation. Following two PBS washes, cultures were serum-starved overnight in basal medium supplemented with 0.1% BSA. On the next day, confluent monolayers were mechanically scratched with a pipette tip to generate a cell-free region. Cells were washed three times with PBS to remove detached cells, treated with the indicated compounds in serum-starvation media and immediately imaged to obtain baseline measurements. The same wound region was imaged again at the indicated experimental endpoint, generally after 1 (PC9, H1975) or 2 (A549) days of treatment. Percentage wound closure was calculated as: Wound closure (%) = [(A₀ − A )/A₀] × 100 where A₀ represents the initial cell-free area and A represents the remaining cell-free area at the endpoint. Because this assay is affected by both proliferation and migration, its endpoint is described as wound repopulation.

### BrdU incorporation and mitomycin C experiments

DNA synthesis was assessed using the Cell Proliferation ELISA, BrdU (colorimetric) kit (11647229001; Sigma-Aldrich) following the manufacturer’s instructions.

BrdU was applied for a 2-h labelling period in serum-starved cultures under the indicated basal or EGF-stimulated conditions. Results were obtained as BrdU incorporation per well.

In separate wound-repopulation experiments, mitomycin C (10 µg/mL; sc-3514B, Santa Cruz Biotechnology) was added 2 h before scratching to restrict proliferation. Mitomycin C was removed by the subsequent PBS washes before treatment initiation.

### Crystal violet staining

Adherent cellular biomass was assessed using crystal violet staining. Cells were fixed with 4% paraformaldehyde for 5 min. Antibody-treated cultures were evaluated at the indicated experimental endpoints, including treatments at 100 nM over 24-72 h (Supplementary Fig. 7). EGFR-silenced and control-siRNA cells were similarly assessed following exposure to the indicated EGFR TKI concentrations (Fig. 8). The dye was solubilized in methanol before absorbance measurement.

### LDH assay

LDH release was measured using the CyQUANT LDH Cytotoxicity Assay (C20301; Thermo Fisher Scientific). At the endpoint of the H1975 wound-repopulation experiments, culture supernatants were collected from the same experimental wells used for wound-closure quantification and analyzed for LDH activity. Three additional wells were treated with lysis buffer to establish the maximum LDH-release reference. Experimental LDH signals were normalized to the mean signal of these three wells, assigned a value of 1.0. Results are expressed as LDH release relative to the maximum-lysis reference rather than percentage cell death.

### EGFR TKI-response experiments

EGFR-silenced and control-siRNA cells were treated with the indicated concentrations of gefitinib or osimertinib, and adherent cellular biomass was assessed using crystal violet staining. Separately, parental LEI and recombinant LEIP were evaluated alone and in combination with gefitinib or osimertinib in H1975 wound-repopulation assays. Supernatants from the same wells were analyzed for LDH activity at the experimental endpoint.

### Three-dimensional PC9 agarose culture

Because PC9 cells formed loose aggregates in ultra-low-attachment plates, an agarose colony-formation approach was used to generate three-dimensional structures.

Cells were dissociated to a single-cell suspension, and 500 cells were plated in each well of a six-well plate in 1.5 mL of complete culture medium containing 0.3% low-melt agarose, layered over 1.5 mL of pre-solidified 0.6% agarose prepared in the same medium. Following solidification of the upper agarose layer, each well was overlaid with 500 µL of culture medium. Cultures were maintained at 37 °C and 5% CO₂ for 14 days, with overlay medium changed every 2-3 days.

The effects of antibody treatment were assessed during formation of the three-dimensional colonies and following their establishment, as indicated in the corresponding experiments.

### H1975 spheroid formation and treatment

H1975 spheroids were generated in ultra-low-attachment Nunclon Sphera plates (174925; Thermo Fisher Scientific) by seeding 1,000 or 5,000 cells per well in 100 µL of culture medium.

Cells were allowed to form spheroids for 48 h before treatment initiation. Spheroids initiated with 1,000 cells per well were used to investigate treatment-associated inhibition of ongoing growth, whereas spheroids initiated with 5,000 cells per well were used to evaluate larger, comparatively stable structures.

On day 0, a partial medium exchange was performed by adding an equal volume of fresh medium, allowing spheroids to settle for at least 30 min and then carefully removing the added volume. Treatments were added in a volume of 20 µL. The partial medium exchange and treatment procedure was repeated over the following two days. Images were acquired on day 0 and day 3.

Changes in projected spheroid area were calculated relative to the initial measurement for each spheroid. Negative values indicate a reduction in projected area relative to baseline.

### Nuclear Green DCS1 imaging

H1975 spheroids were examined using the membrane-impermeant fluorescent DNA dye Nuclear Green DCS1 (catalog 17550; AAT Bioquest) to visualize DNA-associated fluorescence under the indicated treatment conditions. Nuclear Green DCS1 was added at a final concentration of 2 µM, followed by incubation for 1 h before fluorescence imaging. Control and treated spheroids were examined for Nuclear Green-positive material within and surrounding the spheroids., with particular attention to potential baseline core necrosis in untreated structures.

### EGFR immunofluorescence in H1975 spheroids

Upon completion of the corresponding treatments, spheroids from the same treatment were transferred in groups of four using wide-bore tips to microcentrifuge tubes and centrifuged at 30 × g for 5 min. Supernatants were removed while retaining approximately 100 µL to minimize accidental aspiration of spheroids. Spheroids were fixed using paraformaldehyde for 5 min, centrifuged and washed twice with PBS. Samples were not permeabilized and were blocked using 3% BSA for 1 h and incubated overnight with the anti EGFR primary antibody mouse clone 199.12. On the next day, spheroids were incubated with donkey anti mouse AlexaFluor 594 and nuclei were stained with Hoechst 33342. EGFR fluorescence was background-corrected. Binding of LEI to cell-surface EGFR did not interfere with detection by mouse anti-EGFR clone 199.12 (Supplementary Fig. 10).

### Statistical analysis

Where indicated, treatment effects were assessed using one-way or two-way ANOVA, followed by the reported multiple-comparisons procedures, including Šídák’s or Dunnett’s tests. For the H1975 wound-repopulation and LDH analyses, comparisons between antibody treatments within each TKI condition were assessed using Šídák’s multiple-comparisons test, whereas comparisons of TKI-treated groups with the corresponding vehicle condition were assessed using Dunnett’s test. Longitudinal spheroid measurements were analyzed using the statistical procedures indicated in the corresponding figure legends. When measurements were obtained from the same spheroids at baseline and endpoint, the paired structure of the data was considered. Statistical significance was defined at an adjusted P value below 0.05 unless otherwise specified.

## Supplementary figure legends

**Supplementary Figure 1.** Surface-accessible EGFR immunofluorescence in H1975 cells following parental CTX, CTX-R12 or CTX-LINC18 treatment during wound repopulation assay . Green channel - extracellular EGFR domain; Blue channel -nuclei

**Supplementary Figure 2.** C**e**tuximab**, LEI and LEIP prevent EGF-induced EGFR phosphorylation.** Acute EGF-induced EGFR phosphorylation following pretreatment with parental CTX(C) or LEI (L) (left), or following pretreatment with LEI or LEIP (right). Serum-starved A549 cells pretreated with C, LEI or LEIP for 1h before the 5min or 10min (left), or 10 min (right) EGF stimulation at 50ng/ml. Ab4 denominates anti EGFR antibody candidate which was not used in this work.

**Supplementary Figure 3.** Chemical conjugation of R12 or LINC18, but not p56, enhances the functional activity of the independent anti-EGFR antibody LEI . Points represent individual Wells (3-6 replicates) per condition; Bars show mean ± SD. Group means were compared using Welch’s one-way ANOVA followed by Dunnett’s T3 multiple-comparisons test for all 36 pairwise comparisons (family-wise α = 0.05). Letters above groups summarize the adjusted comparisons: groups with no letter in common differ significantly (adjusted P < 0.05), whereas groups sharing at least one letter were not significantly different. **, *** Adjusted p-value <0.01, <0.001

**Supplementary Figure 4.** Wound closure in PC9 is reduced both by chemically conjugated to Cetuximab to R12 and Linc18, and by Linc19 genetically fused to Cetuximab

**Supplementary Figure 5.** Wound closure in H1975 is reduced both by chemically conjugated to Cetuximab to R12 and Linc18, and by Linc19 fused to Cetuximab

**Supplementary Figure 6.** Wound closure in A549 is reduced both by chemically conjugated to Cetuximab to R12 and Linc18, and by Linc19 fused to Cetuximab

**Supplementary Figure 7.** Effects of internalizing anti-EGFR constructs on cellular biomass. Crystal violet staining was used to quantify biomass following treatment of (a) H1299 and A549 cells with CTX-R12 (100 nM, 72 h); (b) A549 cells with CTX-R12 or CTX-LINC18 (100 nM, 24 h); and (c) H1975 and (d) PC9 cells with LEI or LEIP (100 nM, 48 h). Values were normalized to the corresponding untreated Control and are presented as mean ± SD. No statistically significant differences were detected between the groups compared (P > 0.05).

**Supplementary Figure 8.** Validation of siRNA-mediated EGFR depletion by fixed-cell ELISA in PC9 and H1975 cells following transfection with non-targeting control or EGFR-targeting siRNA. Cetuximab was used as primary EGFR recognition antibody. Anti-human HRP secondary and TMB substrate were used to develop the signal.

**Supplementary Figure 9.** S**p**ecificity **of live-cell EGFR detection and compatibility with cetuximab binding.** Relative live-cell ELISA signal in wild-type (WT) and EGFR-silenced (siEGFR) A549 cells, measured using mouse anti-EGFR clone 199.12 (MS-396-P1; Epredia) in the absence or presence of a tenfold excess of cetuximab (CTX). Bound detection antibody was detected using an HRP-conjugated anti-mouse secondary antibody and TMB substrate. Signals were normalized to WT cells without CTX (set to 1). EGFR silencing markedly reduced the signal, whereas detection was largely preserved in the presence of CTX, consistent with recognition of an EGFR epitope that does not compete with CTX binding. Data are presented as mean ± SD of four technical replicates.

**Supplementary Figure 10.** C**o**mpatibility **of EGFR detection by clone 199.12 with LEI binding.** Fixed H1975 cells were stained with mouse anti-EGFR clone 199.12 (MS-396-P1; Epredia) alone (upper row) or in combination with 10x higher concentration of LEI (lower row). Columns show nuclear staining (blue), clone 199.12 staining (red) and LEI staining (green). EGFR staining by clone 199.12 remained detectable in the presence of LEI, supporting compatible binding under the conditions tested. Scale bars, 50 µm.

