## Supplementary Figures for "Internalizing-peptide engineering converts blocking anti-EGFR antibodies into EGFR lysosomal targeting degraders"

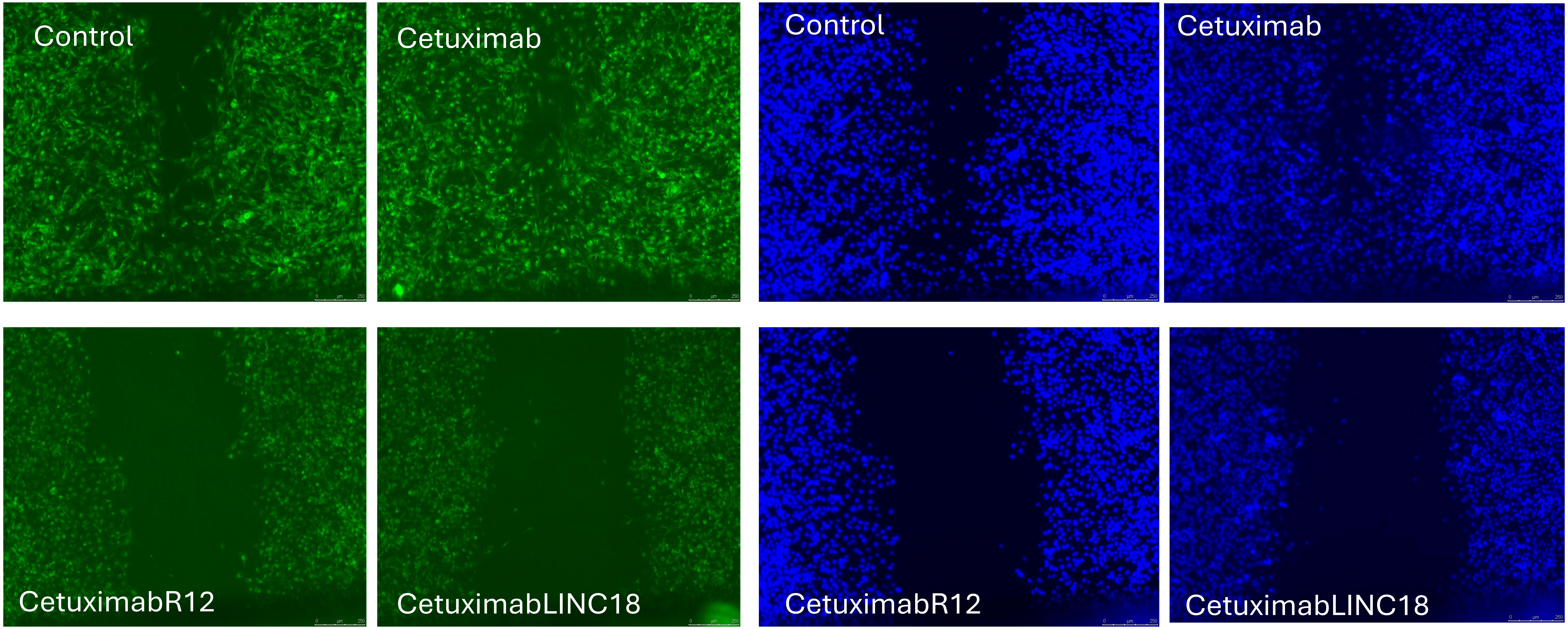


**Supplementary Figure 1. Surface-accessible EGFR immunofluorescence in H1975 cells following parental CTX, CTX-R12 or CTX-LINC18 treatment during wound repopulation assay .** Green channel – extracellular EGFR domain; Blue channel -nuclei

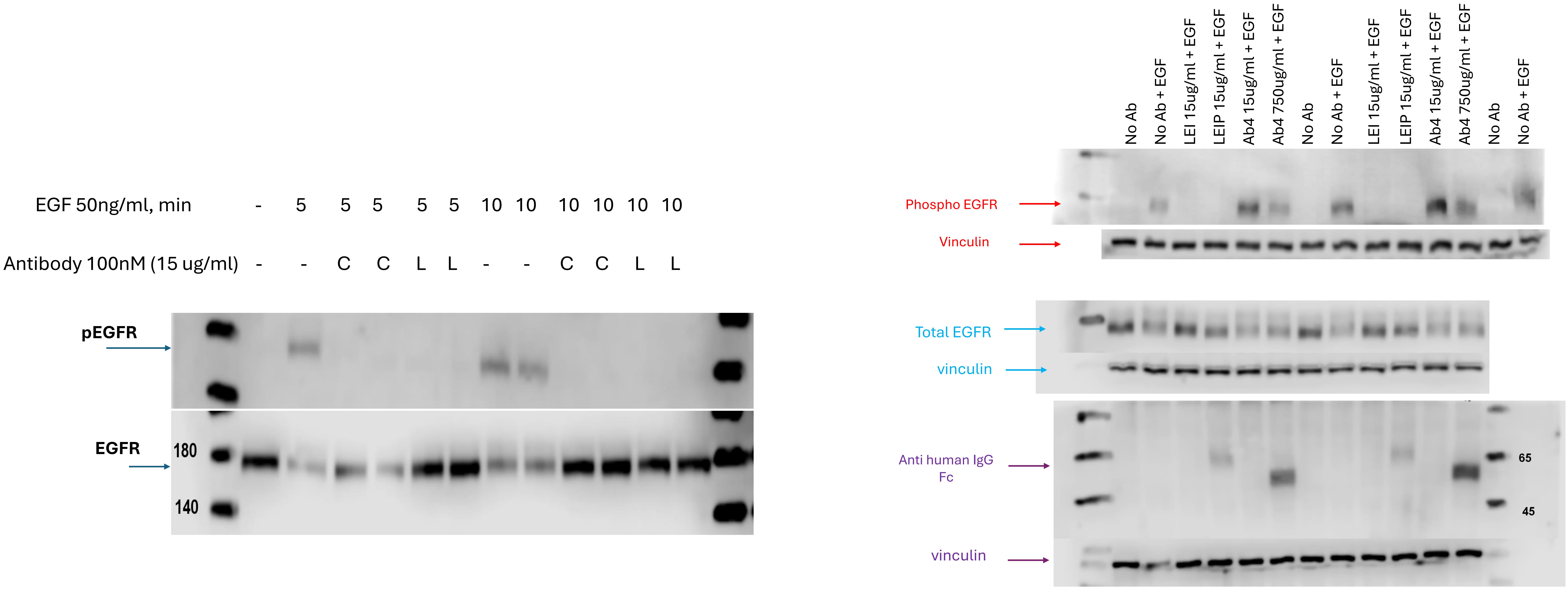


**Supplementary Figure 2. Cetuximab, LEI and LEIP prevent EGF-induced EGFR phosphorylation.** Acute EGF-induced EGFR phosphorylation following pretreatment with parental CTX(C) or LEI (L) (left), or following pretreatment with LEI or LEIP (right). Serum-starved A549 cells pretreated with C, LEI or LEIP for 1h before the 5min or 10min (left), or 10 min (right) EGF stimulation at 50ng/ml. Ab4 denominates anti EGFR antibody candidate which was not used in this work.

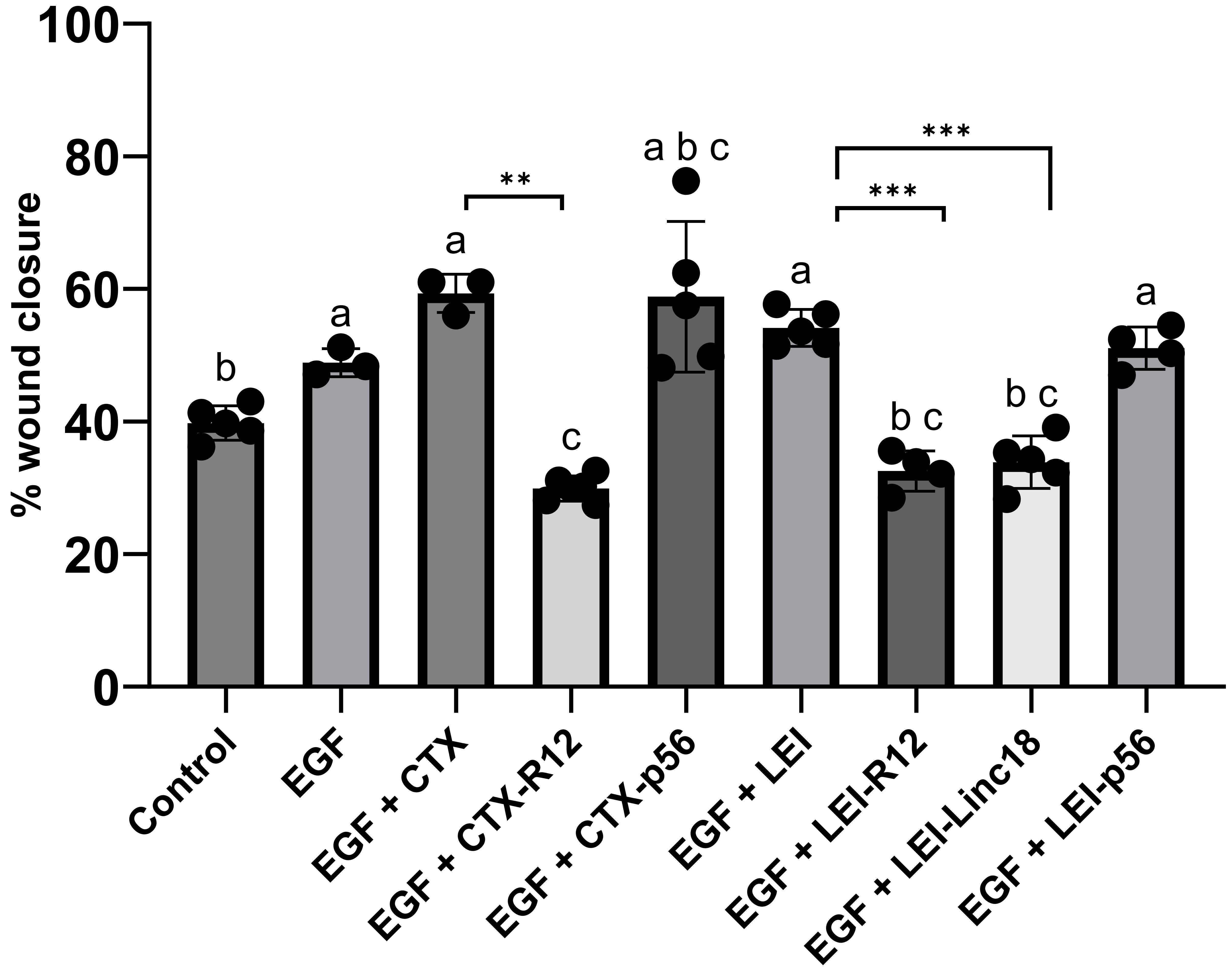


**Supplementary Figure 3. Chemical conjugation of R12 or LINC18, but not p56, enhances the functional activity of the independent anti-EGFR antibody LEI .** Points represent individual Wells (3–6 replicates) per condition; Bars show mean ± SD. Group means were compared using Welch's one-way ANOVA followed by Dunnett's T3 multiple-comparisons test for all 36 pairwise comparisons (family-wise $\alpha = 0.05$). Letters above groups summarize the adjusted comparisons: groups with no letter in common differ significantly (adjusted $P < 0.05$), whereas groups sharing at least one letter were not significantly different. **, *** Adjusted p-value <0.01, <0.001

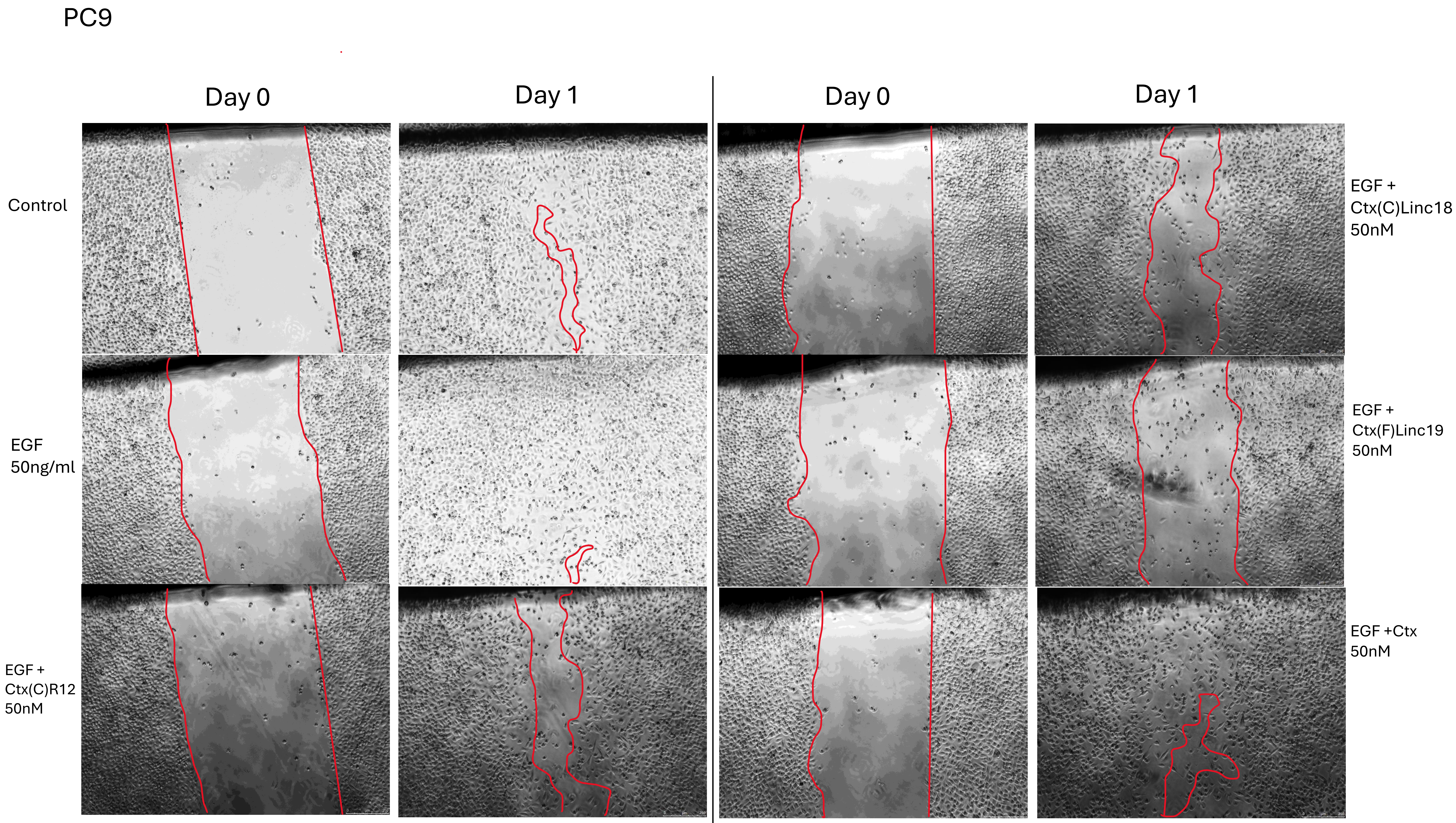


**Supplementary Figure 4** Wound closure in PC9 is reduced both by chemically conjugated to Cetuximab R12 and Linc18, and by Linc19 fused to Cetuximab

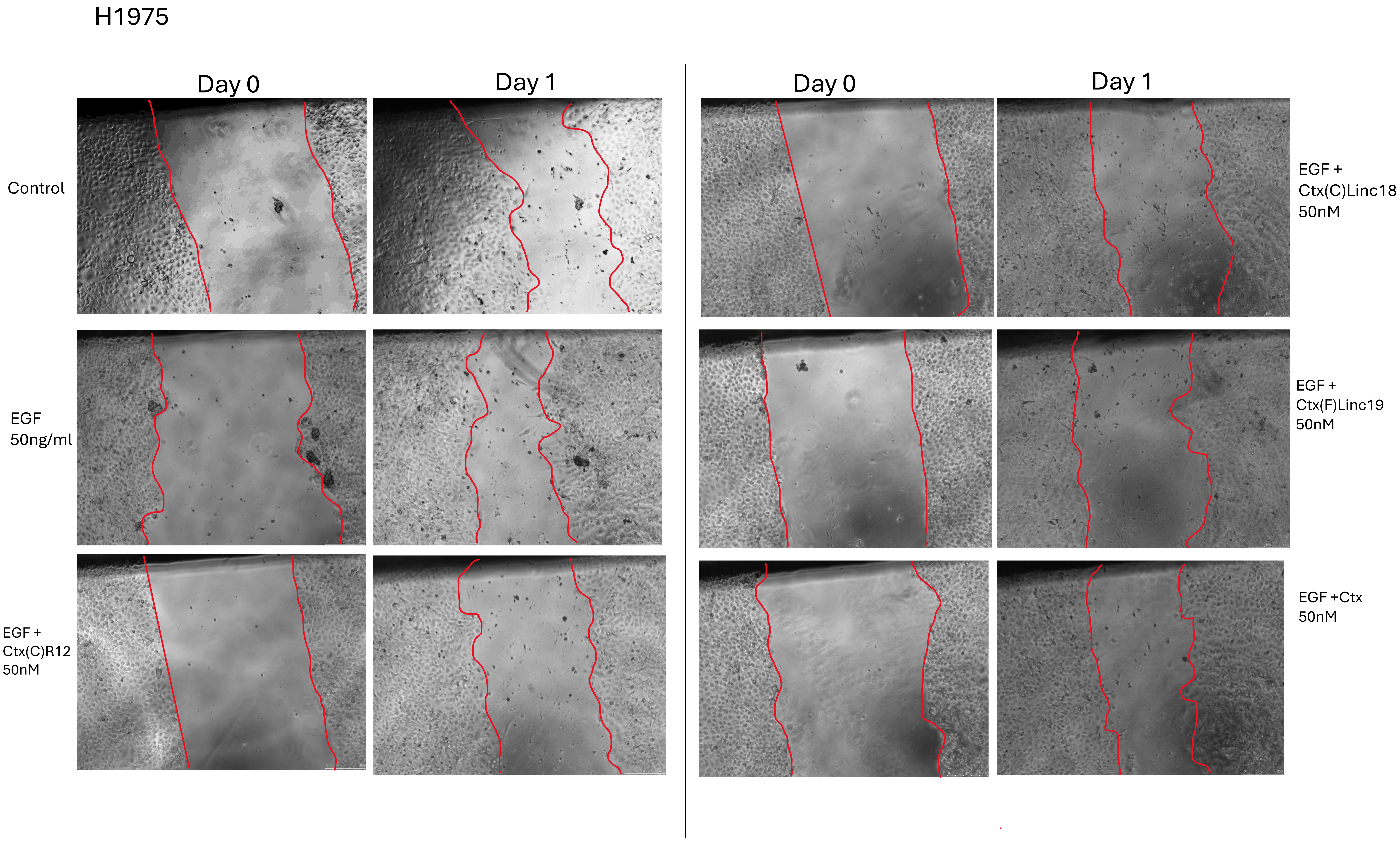


**Supplementary Figure 5** Wound closure in H1975 is reduced both by chemically conjugated to Cetuximab R12 and Linc18, and by Linc19 fused to Cetuximab

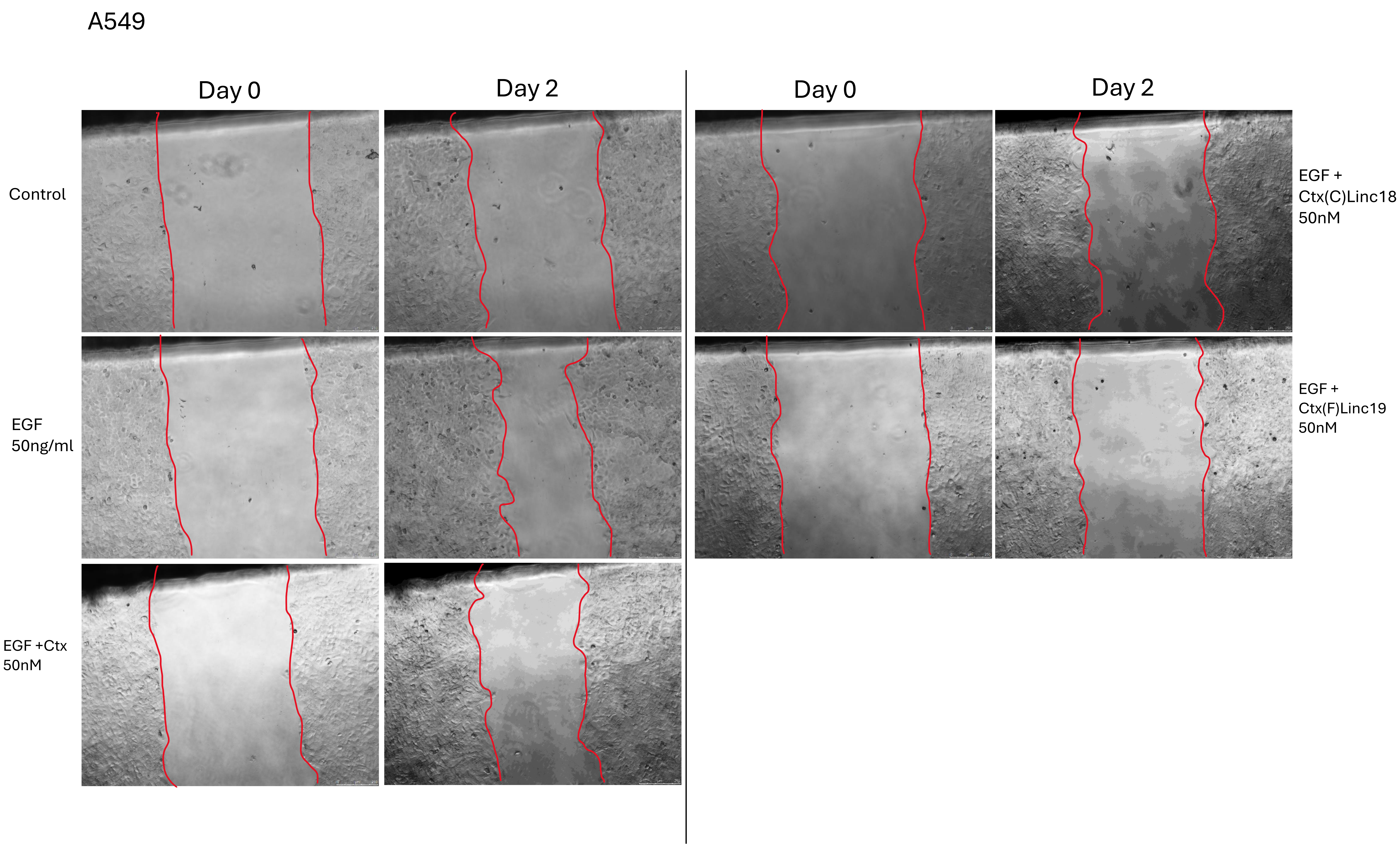


**Supplementary Figure 6** Wound closure in A549 is reduced both by chemically conjugated to Cetuximab R12 and Linc18, and by Linc19 fused to Cetuximab

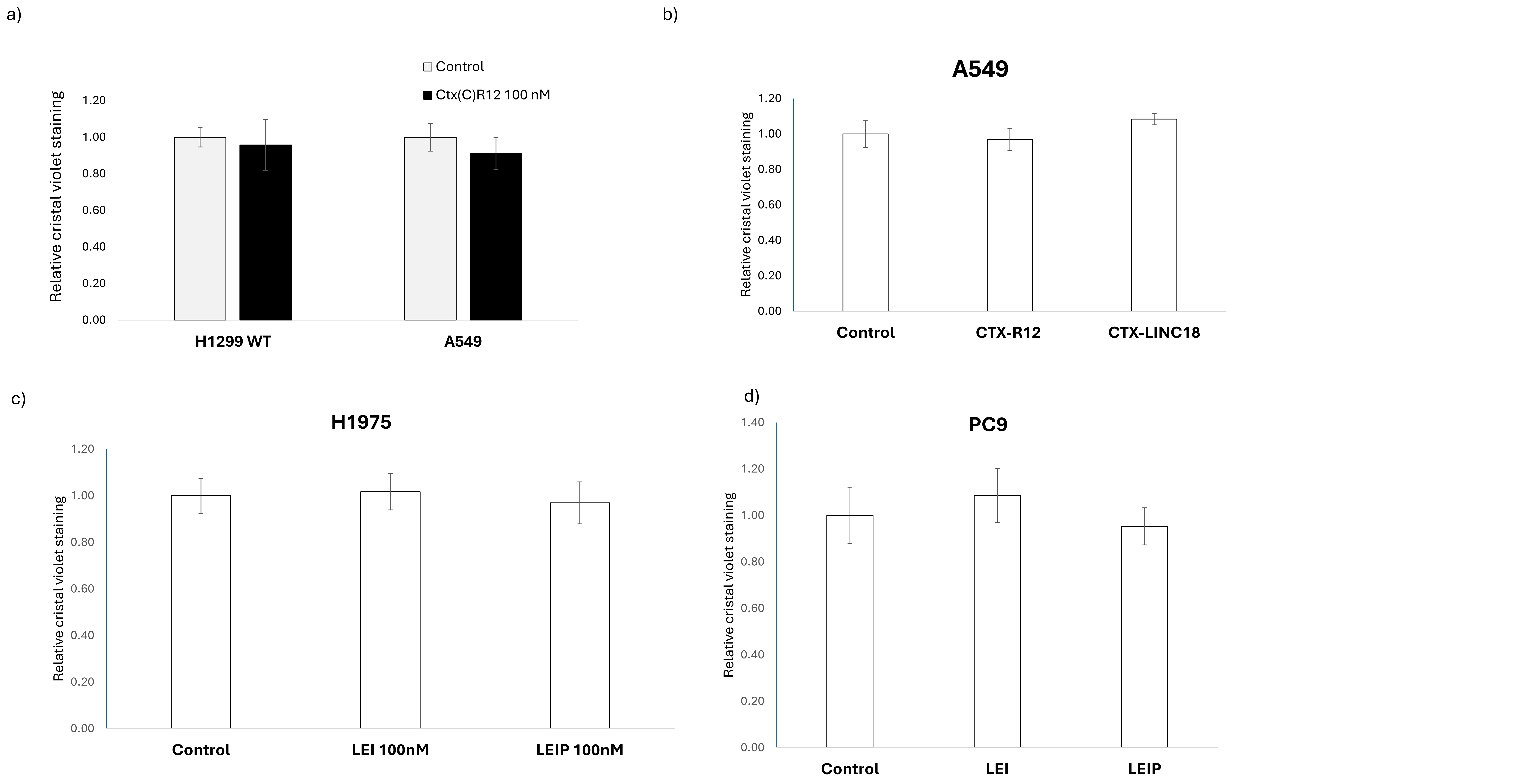


**Supplementary Figure 7. Effects of internalizing anti-EGFR constructs on cellular biomass.** Crystal violet staining was used to quantify biomass following treatment of **(a)** H1299 and A549 cells with CTX-R12 (100 nM, 72 h); **(b)** A549 cells with CTX-R12 or CTX-LINC18 (100 nM, 24 h); and **(c)** H1975 and **(d)** PC9 cells with LEI or LEIP (100 nM, 48 h). Values were normalized to the corresponding untreated Control and are presented as mean ± SD. No statistically significant differences were detected between the groups compared ($P > 0.05$).

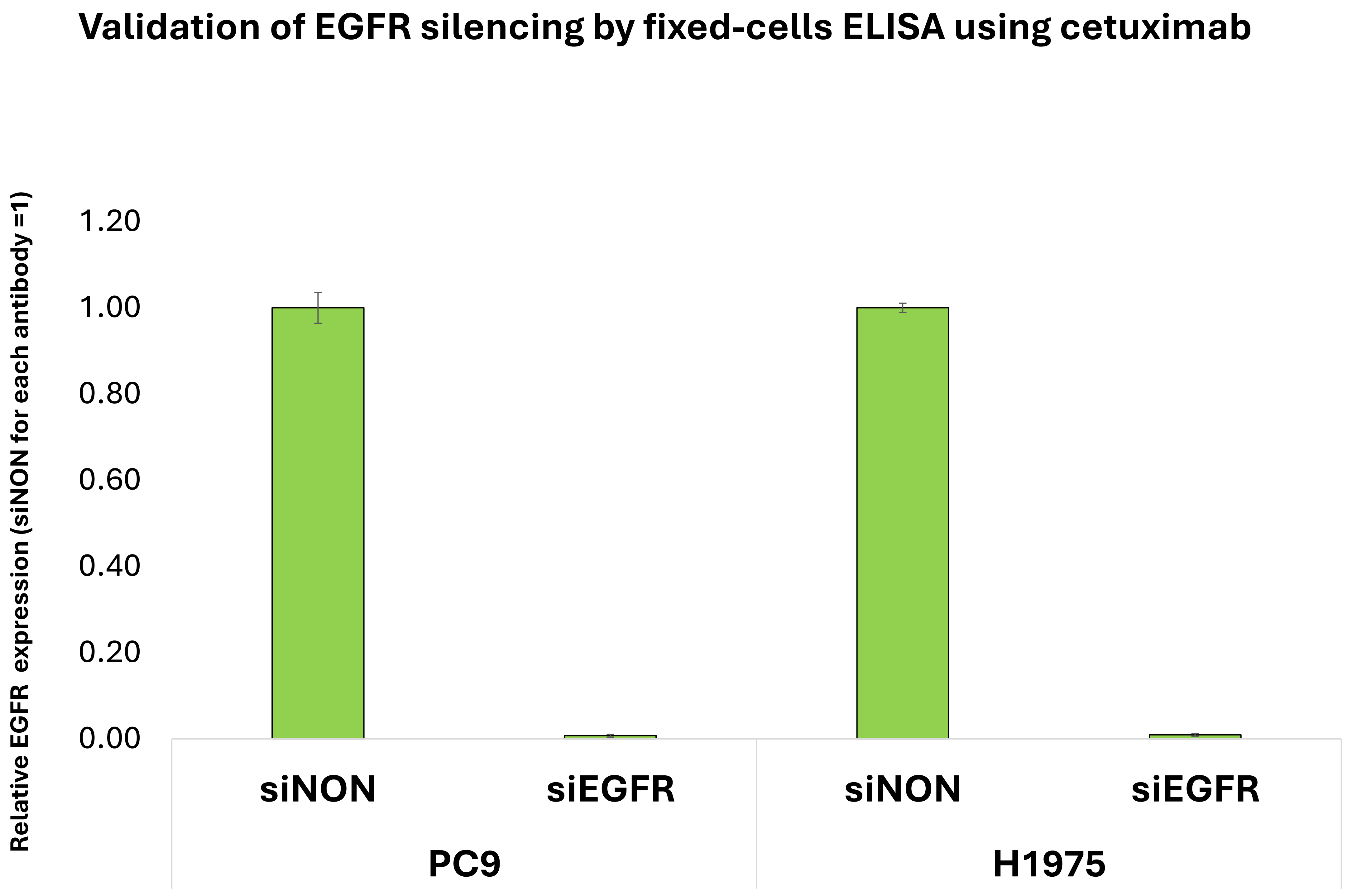


**Supplementary Figure 8.** Validation of siRNA-mediated EGFR depletion by fixed-cell ELISA in PC9 and H1975 cells following transfection with non-targeting control or EGFR-targeting siRNA. Cetuximab was used as primary EGFR recognition antibody. Anti-human HRP secondary and TMB substrate were used to develop the signal.

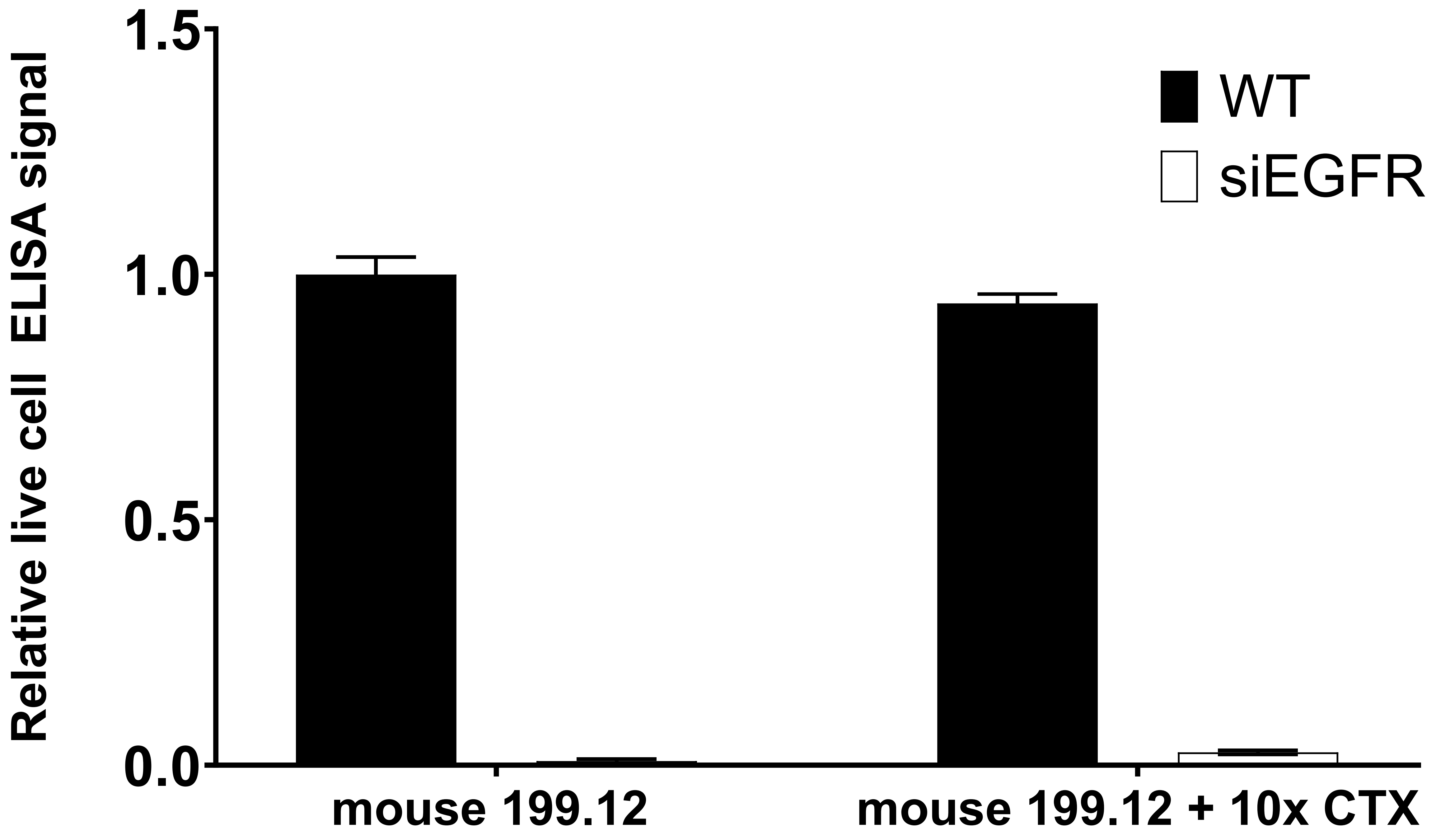


**Supplementary Figure 9. Specificity of live-cell EGFR detection and compatibility with cetuximab binding.** Relative live-cell ELISA signal in wild-type (WT) and EGFR-silenced (siEGFR) A549 cells, measured using mouse anti-EGFR clone 199.12 (MS-396-P1; Epredia) in the absence or presence of a tenfold excess of cetuximab (CTX). Bound detection antibody was detected using an HRP-conjugated anti-mouse secondary antibody and TMB substrate. Signals were normalized to WT cells without CTX (set to 1). EGFR silencing markedly reduced the signal, whereas detection was largely preserved in the presence of CTX, consistent with recognition of an EGFR epitope that does not compete with CTX binding. Data are presented as mean ± SD of four technical replicates.

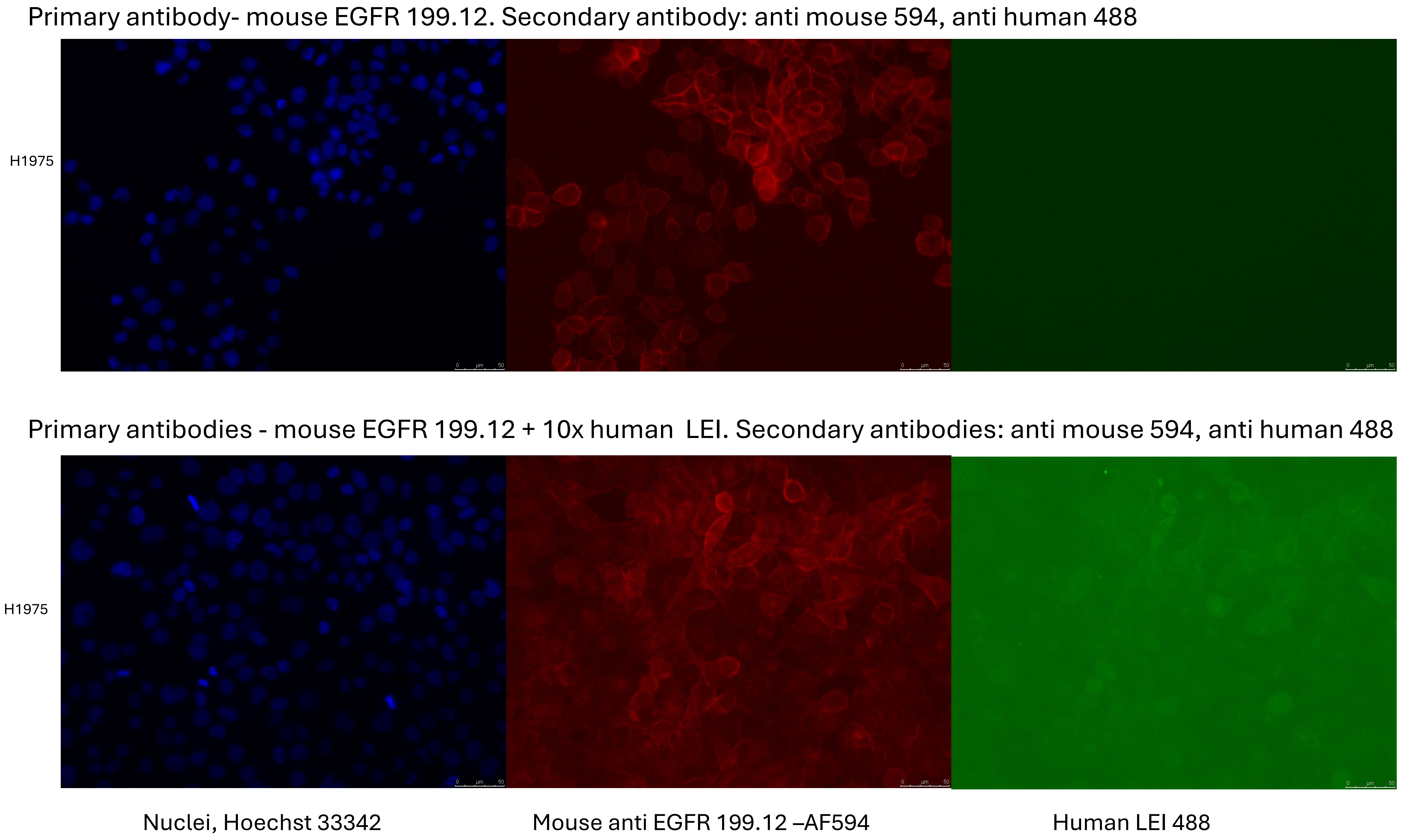


**Supplementary Figure 10. Compatibility of EGFR detection by clone 199.12 with LEI binding.** Fixed H1975 cells were stained with mouse anti-EGFR clone 199.12 (MS-396-P1; Epredia) alone (upper row) or in combination with 10x higher concentration of LEI (lower row). Columns show nuclear staining (blue), clone 199.12 staining (red) and LEI staining (green). EGFR staining by clone 199.12 remained detectable in the presence of LEI, supporting compatible binding under the conditions tested. Scale bars, 50 µm.
